# Modulated termination of non-coding transcription partakes in the regulation of gene expression

**DOI:** 10.1101/2021.01.06.425579

**Authors:** Nouhou Haidara, Odil Porrua

## Abstract

Pervasive transcription is a universal phenomenon leading to the production of a plethora of non-coding RNAs. If left uncontrolled, pervasive transcription can be harmful for genome expression and stability. However, non-coding transcription can also play important regulatory roles, for instance by promoting the repression of specific genes by a mechanism of transcriptional interference. The efficiency of transcription termination can strongly influence the regulatory capacity of non-coding transcription events, yet very little is known about the mechanisms modulating the termination of non-coding transcription in response to environmental cues.

Here, we address this question by investigating the mechanisms that regulate the activity of the main actor in termination of non-coding transcription in budding yeast, the helicase Sen1. We identify a phosphorylation at a conserved threonine of the catalytic domain of Sen1 and we provide evidence that phosphorylation at this site reduces the efficiency of Sen1-mediated termination. Interestingly, we find that this phosphorylation impairs termination at an unannotated non-coding gene, thus repressing the expression of a downstream gene encoding the master regulator of Zn homeostasis, Zap1. Consequently, many additional genes exhibit an expression pattern mimicking conditions of Zn excess, where *ZAP1* is naturally repressed.

Our findings provide a novel paradigm of gene regulatory mechanism relying on the direct modulation of non-coding transcription termination.

## INTRODUCTION

One of the most fascinating discoveries over the last years on the field of gene expression has been the finding that transcription is not restricted to annotated genes but actually extends to the vast majority of the genome. This phenomenon, dubbed pervasive transcription, has been revealed in a multitude of organisms from bacteria to humans (reviewed in 1). Pervasive transcripts are often rapidly degraded, and therefore most of them remain invisible unless RNA degradation is prevented. In general, pervasive transcripts originate from nucleosome-free regions, either from cryptic promoters or from *bona fide* gene promoters that generate transcription divergently from the controlled gene with a certain frequency. Indeed, several studies have concluded that the vast majority of eukaryotic promoters are intrinsically bidirectional (2–5).

Pervasive transcription is potentially detrimental to cell homeostasis because, if left uncontrolled, it can invade and interfere with genomic regions involved in gene regulation, DNA replication or repair (4, 6, 7). Therefore, all organisms studied to date have evolved different mechanisms to limit the negative consequences of pervasive transcription. These mechanisms often rely on early termination of non-coding transcription and ncRNA degradation (reviewed in 1).

The mechanisms responsible for the control of pervasive transcription have been particularly well characterized in budding yeast. In this organism, the so-called NNS (Nrd1-Nab3-Sen1) complex induces early termination of pervasive transcription and promotes the degradation of the resulting ncRNAs by the nuclear exosome (for review, see 8). Nrd1 and Nab3 are RNA-binding proteins that recognize specific sequences that are enriched in the target ncRNAs (4, 9–12), while the conserved helicase Sen1 is the catalytic subunit of the complex (13). The association of Sen1 with Nrd1 and Nab3 enhances Sen1 recruitment to ncRNAs, albeit is not a strict requirement for Sen1 loading on these RNAs (14). Finally, Sen1 uses the energy of ATP hydrolysis to translocate along the nascent RNA and induce the release of RNAPII from the DNA (13, 15).

The widespread character of pervasive transcription has raised the question of its possible biological role. Unlike metazoans, where ncRNAs can play multiple regulatory roles, in budding yeast it is the act of transcription rather than the ncRNA produced which has proved to be functional. Non-coding transcription can interfere with the expression of genes that are located just downstream or in the antisense orientation when it invades the promoter region of these genes. The mechanisms at play typically involve the deposition of repressive chromatin marks at the promoter of regulated genes (16–19). There is a growing number of well-characterized examples where non-coding transcription mediates the regulation of gene expression in response to environmental cues. These include genes involved in amino acid biosynthesis, meiosis, stress, etc (20–22). In many cases, the ncRNA is expressed from a promoter that is only active under specific conditions (i.e. conditions in which the regulated protein-coding gene needs to be repressed). Paradigmatic examples of this mode of regulation are the *ADH1* and *ADH3* genes, which encode two abundant Zn-dependent alcohol dehydrogenases whose expression is repressed in response to Zn limitation by the upstream non-coding genes *ZRR1* and *ZRR2*, respectively. The expression of *ZRR1* and *ZRR2* is activated by the master regulator of Zn homeostasis, Zap1, and transcription through the promoter regions of *ADH1* and *ADH3* interferes with promoter recognition by the corresponding transcriptional activators (23).

More than 1500 non-coding genes expressed under standard growth conditions are targeted by the NNS-dependent pathway for early transcription termination coupled to ncRNA degradation. Importantly, in many of these cases depletion of the Nrd1 component of the NNS-complex abrogates termination by this pathway and provokes misregulation of the neighboring genes (4, 17), indicating that the efficiency of non-coding transcription termination is a potentially relevant parameter in the regulation of gene expression. However, thus far it remains unclear whether the activity of the NNS-complex on non-coding genes can be modulated in response to specific environmental signals to promote gene regulation. To address this question, we have focused on the mechanisms that control the activity of the catalytic subunit of the NNS-complex, the helicase Sen1. We have identified a phosphorylation at a conserved residue of Sen1 catalytic domain and we have shown that phosphorylation at this site (T1623) impairs the interaction of Sen1 with the RNA and reduces the efficiency of transcription termination both *in vitro* and *in vivo*. Genome-wide transcription analyses revealed that a T1623 phospho-mimetic mutation provoked repression of the *ZAP1* gene via transcriptional interference exerted by an upstream non-annotated gene that we have dubbed *ZRN1* for Zap1-Repressor Non-coding gene 1. This resulted in an alteration in the expression of many Zap1-dependent genes involved in Zn homeostasis and an overall transcription profile reminding conditions of Zn excess, where *ZAP1* is naturally repressed. Our data suggested that phosphorylation at Sen1 T1623 could also promote coordinated repression of additional genes by transcriptional interference to further contribute to the regulation of Zn intracellular levels.

Our results support a model whereby the catalytic domain of Sen1 would be phosphorylated under specific conditions to modulate its activity and regulate the expression of genes involved in Zn homeostasis.

## RESULTS

### Identification of a phosphorylation at a conserved threonine of Sen1 helicase domain

To identify possible mechanisms of regulation of Sen1-mediated transcription termination we undertook an analysis of Sen1 post-translational modifications. To this end, we immunoprecipitated a C-terminally TAP-tagged version of Sen1 expressed at the endogenous locus and performed mass spectrometry analyses on Sen1 immunoprecipitates on 5 biological replicates (table S1). We identified 7 phosphosites, among which 6 are novel, spread all along Sen1 functional domains (figure 1A). Sen1 possesses an essential superfamily 1 helicase domain (aa 1095-1905) flanked by a large N-terminal domain (aa 1-975) and a C-terminal intrinsically-disordered region (aa 1906-2231), both involved in protein-protein interactions that are important for Sen1 function (14, 24, 25). We considered particularly interesting the phosphosites located at the helicase domain (HD), since we have previously shown that this domain is necessary and sufficient for transcription termination *in vitro* (15). Among them, phosphorylation of T1623 was detected in two independent experiments and the specific location of this residue made its phosphorylation a good candidate for mediating the regulation of Sen1 catalytic activity. Indeed, a comparison of the structure of Sen1 with that of the closest paralogue of Sen1, Upf1, in complex with the RNA (figure 1B) indicated that T1623 is placed at the predicted RNA-binding region of Sen1. This region could be inferred from the structural similarity between both proteins as well as from our previous mutational analysis confirming that conserved amino acids located at Upf1 RNA-binding surface are also involved in Sen1 interaction with the RNA (26). Because the predicted RNA-binding region of Sen1 is rather positively charged, we hypothesized that phosphorylation of T1623, by introducing a negative charge, could interfere with Sen1 RNA-binding.

**Figure 1:**
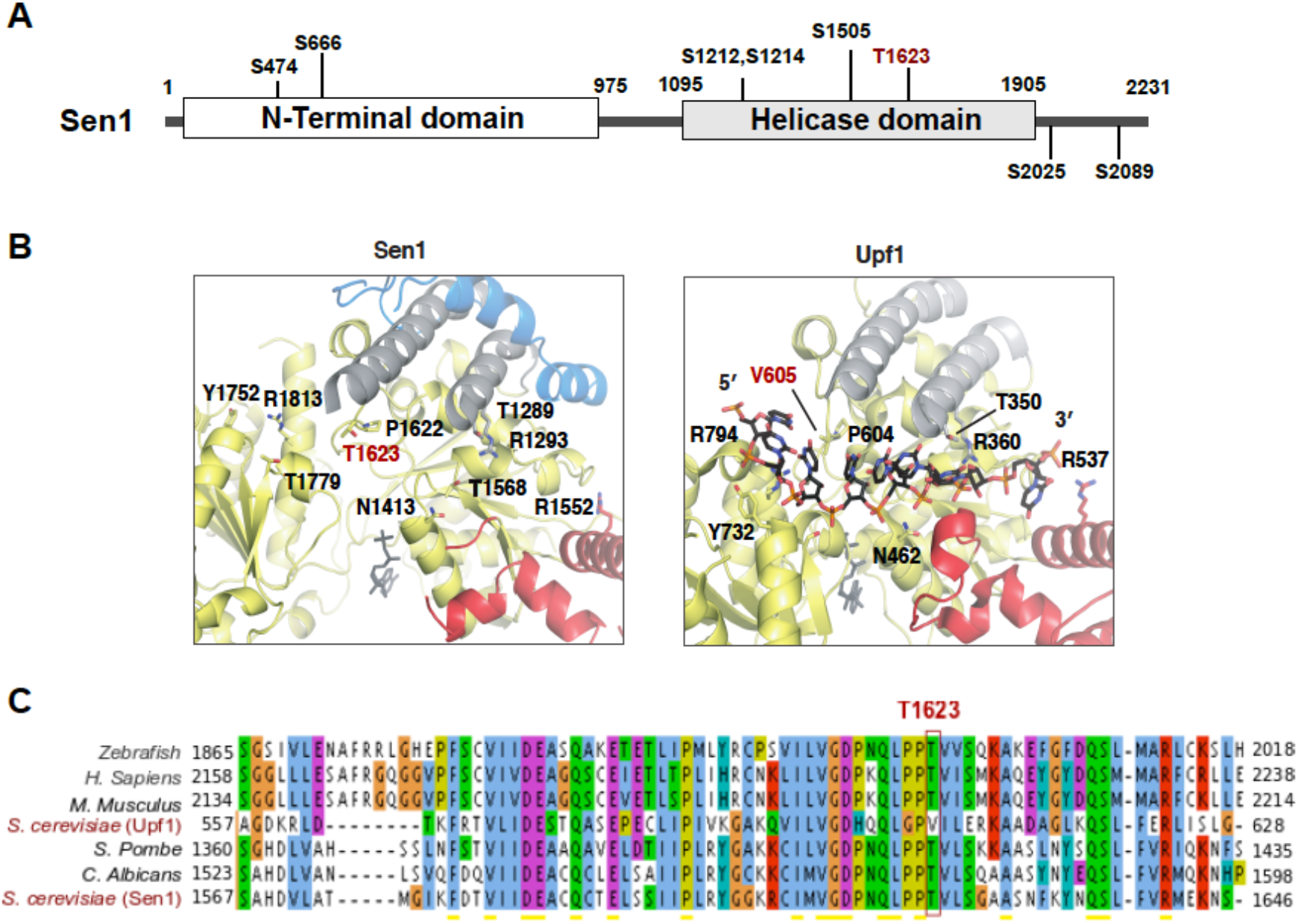
The catalytic domain of Sen1 is phosphorylated at a conserved threonine at the predicted RNA-binding region. **A)** Scheme of Sen1 protein indicating the location of the different phosphosites identified by mass spectrometry. Globular domains are indicated by solid bars whereas regions predicted to be disordered by IUPred (71) are shown by a line. Phosphorylation of residue S1505 was also previously detected in former studies (72, 73). **B)** Zoom in view of the RNA-binding region of Sen1 and Upf1. Protein regions are colored according to the color scheme used for the different subdomains of Sen1 and Upf1 helicase domains in a former study (26). Specifically, amino acids in the core RecA subdomains are indicated in yellow, whereas amino acids in the so-called brace, stalk and prong are shown in blue, grey and red, respectively. Structural data correspond to PDB: 5MZN for Sen1 and PDB: 2XZO for Upf1-RNA complex. Sen1 T1623 and the equivalent V605 in Upf1 are indicated in red. Residues labeled in black are conserved positions involved in RNA binding by Upf1 that revealed a role in Sen1 interaction with RNA in a former mutational analysis (26). **C)** Alignment of a small fragment of Sen1 and Upf1 proteins containing T1623 or the equivalent residues. Amino acids are colored according to the clustal color scheme.

Alignment of Sen1 proteins from diverse organisms revealed that the T1623 residue is highly conserved from budding yeast to humans (figure 1C), further supporting the idea that this threonine is important for Sen1 activity. Interestingly, in Upf1 the equivalent amino acid is a valine (V605, see figure 1B), which differs from a threonine exclusively in the presence of a methyl instead of the hydroxyl group of the threonine. This suggests that a threonine at that particular position might have been selected in Sen1 proteins throughout evolution either to confer specific properties to Sen1 HD or to confer the capacity to modulate Sen1 catalytic activity by phosphorylation.

### Phosphorylation of Sen1 at T1623 partially inhibits Sen1 activity both in vivo and in vitro

In order to assess the role of T1623 phosphorylation in the regulation of Sen1 function, we introduced mutations either to prevent (T to V) or to mimic (T to E) phosphorylation of this residue and we analyzed the effect of these mutations on cell growth (figure 2A). Mutation to valine did not impair cell growth at any of the temperatures tested. In contrast, the T1623E phospho-mimetic mutation induced a strong growth defect at 37°C, while growth at other temperatures was not significantly affected. This thermosensitive phenotype was not due to decreased stability of the protein at high temperature, since Sen1 protein levels remain similar in cells grown at 30°C and 37°C (figure S1). Because termination of non-coding transcription by Sen1 is essential for viability, this results suggests that phosphorylation might affect negatively Sen1 capacity to promote transcription termination. To test this possibility, we analysed by northern blot the expression of two typical Sen1 targets, namely the CUT (cryptic unstable transcript) NEL025c and the snoRNA *SNR13* (figure 2B). CUTs represent one of the major classes of pervasive transcripts while snoRNAs are functional non-coding RNAs that depend on the NNS-pathway for transcription termination and 3’ end processing (27). Importantly, the efficiency of transcription termination at both non-coding genes was identical in the *sen1T1623V* mutant compared to the wild-type (wt) strain, indicating that a T and a V are interchangeable at position 1623 for normal Sen1 function in transcription termination. On the contrary, in the *sen1T1623E* mutant we observed a mild accumulation of transcripts ∼200-500 nt longer than in the wt, which is suggestive of delayed transcription termination. This accumulation was very moderate under permissive temperature (i.e. at 30°C) but was strongly exacerbated in cells grown for 2h at 37°C (figure S1B), consistent with the thermosensitive phenotype of the *sen1T1623E* mutant. Taken together, these results strongly suggest that phosphorylation of Sen1 T1623 modulates negatively the catalytic activity of Sen1 and, therefore, the efficiency of Sen1-dependent transcription termination.

**Figure 2:**
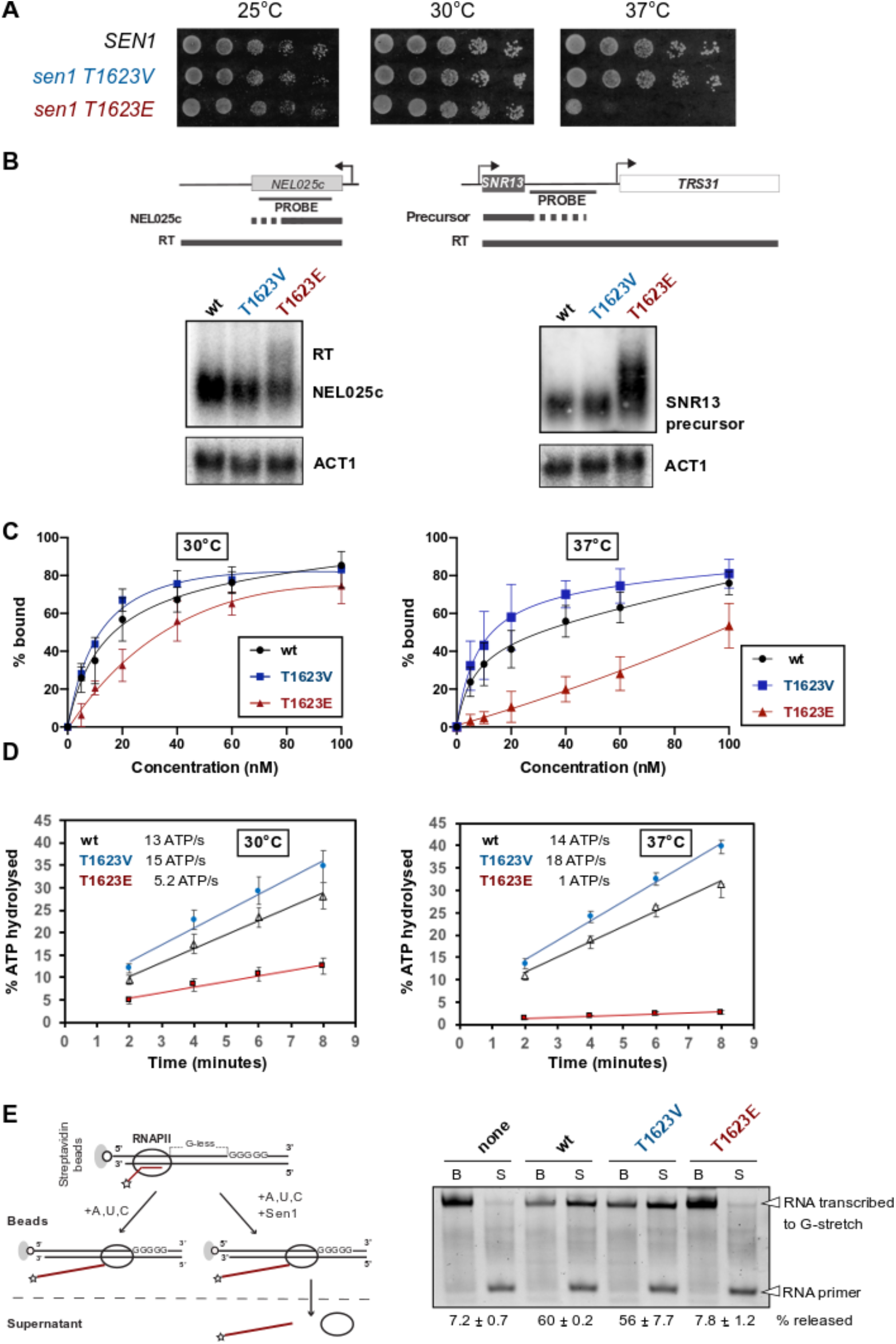
Functional analyses of mutations preventing (T to V) or mimicking (T to E) phosphorylation of Sen1 T1623. **A)** Growth tests of yeast cells expressing different *SEN1* variants on YPD medium at the indicated temperatures. **B)** Northern blot analysis of two typical Sen1 targets in strains expressing either the wt, the T1623V or the T1623E version of *SEN1*. Cells were grown at 30°C. Experiments were performed in a *¹¹rrp6* background to stabilize and, thus, render detectable the primary products of termination by Sen1. The *ACT1* mRNA is used as a loading control. Probes used for RNA detection are described in table S7. **C)** Electrophoretic mobility shift assay (EMSA) using a 5′-end fluorescently labeled 40-mer RNA as the substrate (DL3508, see table S7) at 4 nM and recombinant Sen1 HD variants at 5, 10, 20, 40, 60, and 100 nM at the final concentrations. The plots correspond to the quantifications of three independent experiments performed for each incubation temperature. The values indicate the mean and standard deviation (SD). The dissociation constant (Kd) at 30°C is 13.1 nM for the wt, 12.7 nM for the T1623V and 100 nM for the T1623E version of Sen1 HD. At 37°C the Kds are 6.4 nM, 8.1 nM and > 5μM for the wt, T1623V and T1623E, respectively. Representative gels of these experiments are shown in figure S2B. **D)** Time course analysis of the nucleic acid-dependent ATPase activity of the indicated Sen1 HD variants at either 30°C or 37°C. Values represent the average and SD from three independent experiments. **E)** *In vitro* transcription termination (IVTT) assays with the indicated versions of Sen1 HD Left: Scheme of an IVTT assay. Ternary elongation complexes (ECs) composed of purified RNAPII, fluorescently labeled nascent RNA, and DNA templates are assembled and attached to streptavidin beads via the 5′ biotin of the non-template strand to allow subsequent separation of beads-associated (B) and supernatant (S) fractions. A star denotes the presence of a FAM at the 5′ end of the RNA. The transcription template contains a G-less cassette followed by a G-stretch in the non-template strand. After addition of a mixture of ATP, UTP and CTP at a final concentration of 1 mM each, RNAPII transcribes until it encounters the G-stretch. Sen1 HD dissociates ECs stalled at the G-stretch, therefore releasing RNAPIIs and the associated transcripts to the supernatant. The fraction of transcripts released from ECs stalled at the G-stretch is used as a measure of the efficiency of termination. Right: Representative gel of one out of two independent IVTT assays. The values on the bottom correspond to the average and the SD of three independent experiments.

To gain mechanistic insight into the impact of T1623 phosphorylation on Sen1 catalytic activity, we expressed and purified from *E. coli* the wt, the T1623V and the T1623E versions of Sen1 HD (figure S2) and we characterized them with various biochemical assays. As indicated above, the HD of Sen1 recapitulates *in vitro* the mechanisms of termination employed by the full-length protein (15, 26). These mechanisms involve the interaction of Sen1 with the nascent transcript and Sen1 translocation along the RNA towards RNAPII, which strictly requires ATP-hydrolysis by Sen1 (13, 15, 26). To assess whether phosphorylation of T1623 would affect the association of Sen1 with the RNA, as the location of this residue predicts, we performed electrophoretic mobility shift assays (EMSA) with the three purified versions of Sen1 HD and a fluorescently-labeled 40-mer RNA as the substrate (figure 2C and S2). These experiments were carried out at 30°C and 37°C. The presence of the T1623V mutation did not decrease the affinity of Sen1 for the RNA at any of the temperatures tested. However, we observed a mild decrease in the affinity of Sen1T1623E for the RNA compared to the wt at 30°C, which was significantly more pronounced at 37°C. Because the interaction of Sen1 with the RNA is required to activate Sen1 ATPase activity, we performed ATP hydrolysis assays using non-saturating concentrations of RNA to assess the RNA-dependent ATPase activity of the different Sen1 HD variants (figure 2D). Consistent with the results of EMSA experiments, we observed reduced levels of ATP hydrolysis by the T1623E but not the T1623V version of Sen1, compared to the wt at 30°C. Furthermore, this decrease in Sen1 ATPase activity was strongly exacerbated at 37°C. Taken together, these results strongly suggest that the phosphorylation of Sen1 T1623 destabilizes the interaction of Sen1 with the RNA and provide the molecular basis of the thermosensitive phenotype observed *in vivo*.

To assess whether impaired RNA-binding in the T1623E mutant results in decreased transcription termination efficiency *in vitro*, we employed a highly purified *in vitro* transcription termination (IVTT) assay that we have previously developed (28, see methods for details). In this system, we employ purified RNAPII, transcription templates and Sen1 HD and we monitor the capacity of the different versions of Sen1 to promote the release of paused RNAPIIs and associated transcripts from the DNA template (figure 2E). In agreement with the former results, the T1623V variant behaved as the wt in IVTT assays. In contrast, the T1623E mutant was dramatically affected in its capacity to dissociate the RNAPII from the DNA, both at 30°C and 37°C (figure 2E and S2). Although the *in vitro* system seems more sensitive than the *in vivo* conditions to Sen1 dysfunction, these results confirm that the T1623E phospho-mimetic mutation impairs Sen1 transcription termination activity.

Overall, the results of our *in vitro* biochemical assays provide strong support to the notion that phosphorylation of Sen1 T1623 modulates negatively the capacity of Sen1 to induce transcription termination by interfering with its association with the RNA. Furthermore, they suggest that a threonine rather than a valine at that particular position of the helicase domain might have been selected in Sen1 proteins to provide the capacity to regulate their catalytic activity by phosphorylation.

### Phosphorylation of Sen1 T1623 causes deregulation of a small subset of functionally-related protein-coding genes

The identification of Sen1 T1623 phosphorylation as a mechanism of regulation of Sen1-mediated transcription termination prompted us to search for the targets of such regulation. To this end, we performed a genome-wide analysis of transcription in the presence of the T1623E phospho-mimetic mutation, hoping that this mutation would be sufficient to reproduce specific gene expression patterns that could inform about the regulatory function of Sen1 phosphorylation. Thus, we generated high-resolution maps of transcribing RNAPII by UV crosslinking and analysis of cDNA (CRAC, 29) in yeast strains expressing either the wt or the T1623E version of *SEN1* (figure 3). The CRAC technique relies on the detection and quantification of nascent transcripts that crosslink to and co-purify with transcribing RNAPII. Experiments were performed on two independent biological replicates that yielded very reproducible results (figure S3). We first analyzed the distribution of RNAPII at non-coding genes whose transcription is normally terminated by Sen1 (i.e. snoRNA genes and CUT genes) to assess to what extent the transcription termination defects observed at individual targets in the T1623E mutant are widespread. Metagene analyses of snoRNA genes revealed a moderate increase in the RNAPII signal downstream of the annotated transcription termination site, indicating mild transcription termination defects (figure 3A). More detailed heatmap analyses of individual snoRNA genes showed that these termination defects applied to most snoRNA genes and consisted in a delay in termination of ∼300 nt on average (figure 3B and representative example in figure 3C). A similar behaviour was observed at CUTs, although the termination defects were less pronounced than at snoRNA genes (figure 3D-F). Altogether, these results indicate that the T1623E mutation provokes moderate transcription termination defects at a majority of, albeit not all, non-coding genes that depend on Sen1 for termination.

**Figure 3:**
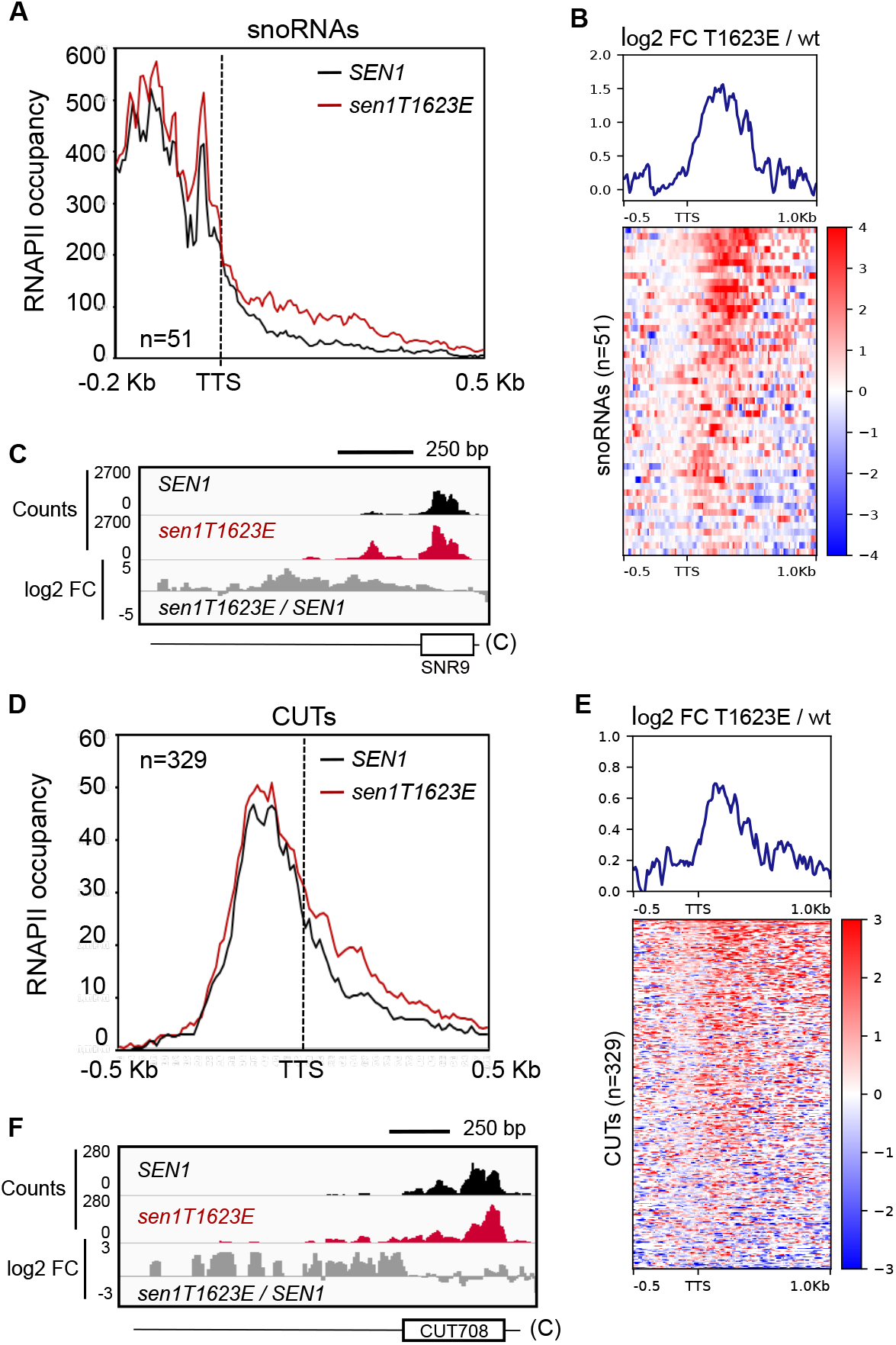
A phospho-mimetic mutation in Sen1 T1623 globally reduces the efficiency of transcription termination at non-coding genes. **A)** Metagene analysis of the RNAPII distribution relative to the annotated transcription termination site (TTS) of snoRNAs. Values on the *y*-axis correspond to the median coverage. **B)**Heatmap analysis representing the log2 of the fold change (FC) of the RNAPII signal in the *sen1T1623E* mutant relative to the wt at snoRNA genes. The summary plot on the top was calculated using the average values for each position. **C)** Integrative Genomics Viewer (IGV) screenshots of an example of snoRNA genes displaying termination defects in the *sen1T1623E* mutant. The values correspond to the counts per million of counts (CPM) multiplied by 10. **D)** Metagene analyses of transcribing RNAPII at CUTs performed as in **A**). **E)** Heatmap analyses of CUTs performed as in **B**). **F)** IGV screenshot of an example of CUTs exhibiting transcription termination defects in the *sen1T1623E* mutant.

We next performed a differential expression analysis on protein-coding genes, that we expected to be the ultimate targets of regulation by Sen1 T1623 phosphorylation (figure 4A). Interestingly, despite the relatively widespread nature of the termination defects observed in the *sen1T1623E* mutant, our analyses revealed significant changes in the expression of only a small (∼1%) subset of protein-coding genes. Specifically, we detected 66 genes that were upregulated and 53 genes that were downregulated in the *sen1T1623E* mutant. We manually curated this gene list to exclude misannotated genes and others false positives (see methods for details) and we ended up with a more robust set of 84 genes that was subjected to more in depth analyses (table S2).

**Figure 4:**
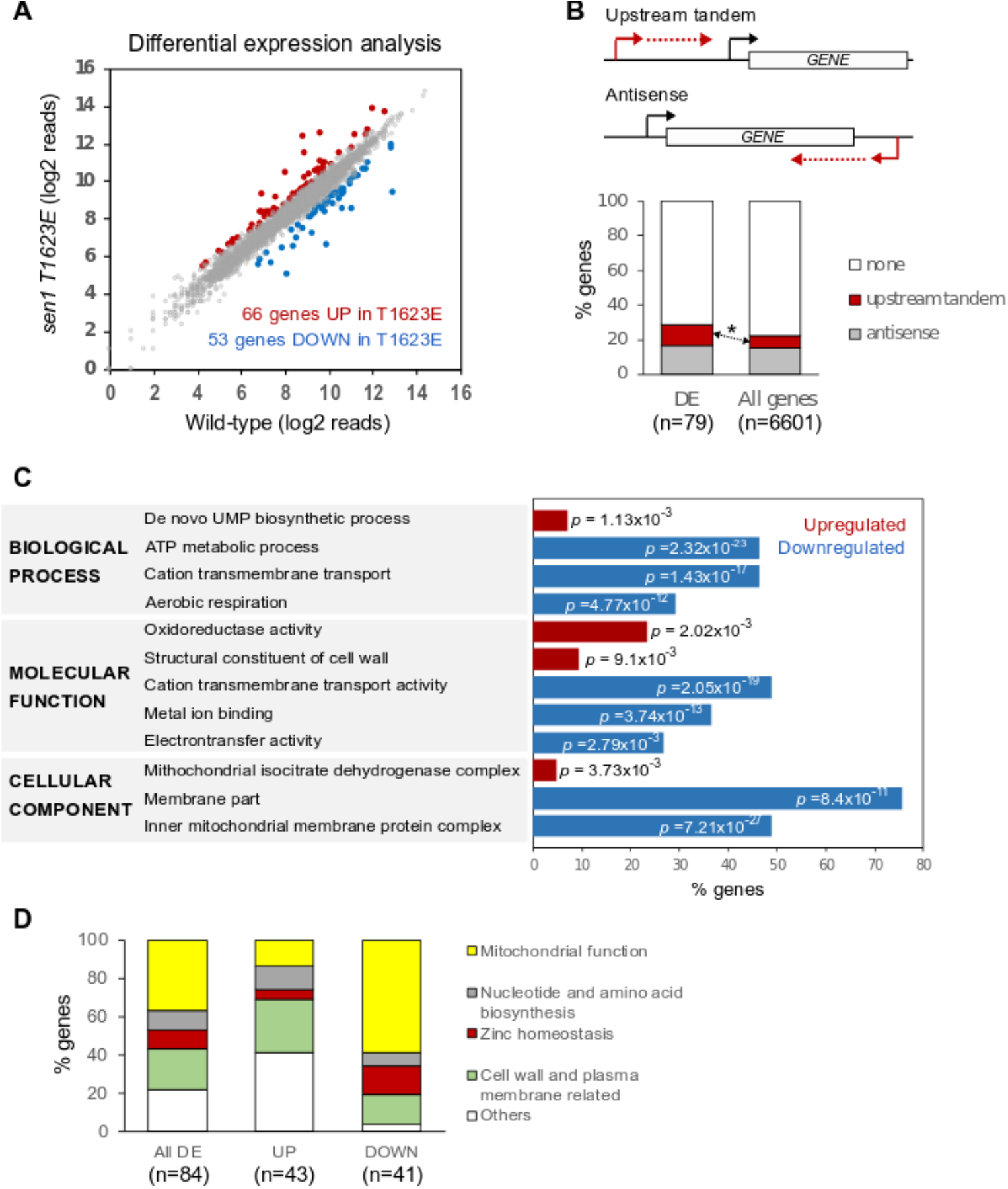
The T1623E mutation in Sen1 induces changes in the expression of functionally-related genes. **A)** Analysis of differentially expressed protein-coding genes in the *sen1T1623E* mutant relative to the wt. Genes are considered as upregulated or downregulated if the log2 fold change (FC) *sen1T1623E / wt is >0*.*5* or *<-0*.*5*, respectively, and the p-value <0.05. The full list of genes with their corresponding expression levels and associated p-values is provided in table S2. **B)**Analysis of non-coding transcription relative to genes differentially expressed (DE) in the *sen1T1623E* mutant relative to the wt. An asterisk denotes a statistically significant (p-value = 0.02) enrichment in genes transcribed immediately downstream of annotated CUTs and SUTs. Genes encoded in the mitochondrial genome were excluded from these analyses since they are transcribed by a dedicated RNAP. **C)** Representative subset of gene ontology (GO) terms found significantly overrepresented in in the *sen1T1623E* mutant relative to the wt. *p* represents the associated p-value. The full list of GO terms is provided in tables S3 and S4.

To explore the connections between the defects in non-coding transcription termination and the changes in the expression of protein-coding genes provoked by the Sen1 T1623E mutation, we performed an analysis of non-coding transcription relative to differentially-expressed (DE) protein-coding genes (figure 4B). Because genes that are regulated by non-coding transcription are typically located either downstream of or antisense to non-coding transcription units, we reasoned that genes in either or both configurations might be overrepresented in the set of DE genes compared to the whole protein-coding genome. Indeed, we observed a moderate but statistically significant enrichment in genes located just downstream of annotated non-coding genes (CUTs and Stable Unannotated Transcripts or SUTs, 5) whose transcription is typically terminated by the NNS-dependent pathway (4). This suggests that at least part of the changes in gene expression we observed in the *sen1T1623E* mutant could be due to events of transcriptional interference by impaired termination of non-coding transcription just upstream of the repressed genes.

Next we performed a gene ontology analysis on the set of DE genes to search for possible signatures or functional connections between these genes. Interestingly, several common themes emerged, especially from the set of downregulated genes (figure 4C and tables S3 and S4). Indeed, GO terms related to transport, energetic metabolism and mitochondrial activity were strongly overrepresented in the list of downregulated genes. Furthermore, regarding the cellular location of the encoded proteins, the vast majority of downregulated genes corresponded to cell wall and membrane proteins and an important fraction of them were actually part of the inner mitochondrial membrane. Concerning the upregulated genes, we found a modest enrichment in genes involved in UMP synthesis, likely related to the effects of the T1623E mutation on *URA2* (figure S4), a well-characterized gene regulated by Sen1-mediated transcription termination (30). We also detected an enrichment in genes encoding cell wall and mitochondrial components.

Guided by the results of our GO analysis we investigated the function of each DE gene individually and we classified genes into non-overlapping functional categories that could facilitate subsequent analyses (figure 4D and table 1). In this way we found that roughly 40% of DE genes encoded proteins related to mitochondrial functions, about 20% of genes were assigned to cell wall and plasma membrane related functions and ∼10% participated in nucleotide and amino acid biosynthetic pathways. Interestingly, we also found that ∼10% of DE genes were involved in Zn homeostasis. Finally, about 20% of genes, grouped as “others”, corresponded mainly to genes with unknown functions and genes encoding transcription factors or vacuolar proteins. When we inspected separately upregulated and downregulated genes, we found that genes involved in mitochondrial functions and Zn homeostasis were mostly downregulated, while the other two main functional groups were composed of more similar numbers of up-Sand downregulated genes.

**Table 1:** List of genes differentially expressed in the *sen1T1623E* mutant grouped by functional categories. Upregulated genes are indicated in red, whereas downregulated genes are coloured in blue.

| Functional category | Genes |
| --- | --- |
| Mitochondrial-related | *IDH1*, *IDH2*, *COX2*, *COX3*, *DLD1*, *CIS1*, *ACO1*, *ATP9*,<br>*INH1*, *COX1*, *COB*, *ATP7*, *COX4*, *ATP1*, *ATP2*, *ATP3v*,<br>*ATP5*, *PSD1*, *COR1*, *QCR2*, *MDH1*, *ATP4*, *ATP16*,<br>*COX8*, *COX6*, *QCR9*, *RIP1*, *SDH2*, *MIN8*, *SDH1*, *ATP6* |
| Nucleotide and amino acid biosynthesis | *IMD2*, *ARG3*, *URA2*, *URA1*, *URA3*, *LEU1*, *LEU2*, *MET6* |
| Zinc homeostasis | *ADH1*, *ADH2*, *ZRT1*, *ADH4*, *ZPS1*, *ZAP1*, *MNT2*, *ZRT3* |
| Cell wall and plasma membrane-related | *PDR5*, *AQR1*, *SPO24*, *PRM5*, *YEH1*, *KTR2*, *NCW2*,<br>*PIR3*, *YPS3*, *PDR16*, *CRH1*, *SED1*, *PHO84*, *SAM2*,<br>*HXT4*, *SAM3*, *ITR1*, *TPO4* |
| Other functions | *PIR5*, *YGR035C*, *MDH2*, *SHU1*, *YCL049C*, *VID24*, *SCE1*,<br>*TRM13*, *ATG40*, *SRD1*, *RRT15*, *HRP1*, *ACA1*, *HTC1*,<br>*MIG3*, *CRG1*, *DPI8*, *INO1*, *VTC3* |

In summary, these results suggest that phosphorylation of Sen1 T1623 could play a role in the regulation of specific pathways and cellular processes.

### The T1623 phospho-mimetic mutation provokes repression of the gene encoding the master regulator of Zn homeostasis by transcriptional interference

We decided to focus on genes related to Zn homeostasis since, as mentioned above, several genes involved in this process were previously shown to be regulated by non-coding transcription (23). Interestingly, when we examined carefully the genes from this group, we found that the gene encoding the master regulator of Zn homeostasis, Zap1, is located just downstream of a non-annotated transcription unit (figure 5A). The transcripts produced by this putative non-coding gene behave as CUTs because RNAseq data revealed that these RNAs are strongly stabilized by deletion of *RRP6*, which encodes one of the major exonucleases responsible for degradation of CUTs (2, 5, 31). Importantly, the Sen1 T1623E phospho-mimetic mutation induced transcription termination defects at this non-coding gene, which resulted in increased transcription through the promoter region of *ZAP1* and, consequently, decreased RNAPII signal at *ZAP1* gene body and decreased levels of *ZAP1* mRNA (figure S5). These observations strongly suggest that phosphorylation of Sen1 T1623 promotes repression of the *ZAP1* gene by a mechanism of transcriptional interference mediated by impaired transcription termination of an upstream non-coding gene. We dubbed this novel non-coding gene *ZRN1* for Zap1-Repressor Non-coding gene 1 and we used our CRAC datasets together with former annotations of transcriptional start sites in *!ιrrp6* (32) to define the coordinates of this gene as positions 334,108 (start) and 333,439 (end) of chromosome X. Zap1 is a transcriptional activator that recognizes a sequence motif called Zn Responsive Elements (ZRE) at the promoter region of the genes it regulates and recruits the SWI/SNF, SAGA, and Mediator complexes to promote the expression of genes involved in Zn uptake, Zn mobilization from vacuolar stores, etc (33). Zap1 regulates its own expression by binding to a ZRE that is located at the region between *ZRN1* and *ZAP1* genes, suggesting that increased transcription through this region in the *sen1T1623E* mutant could impair the recognition of this sequence by Zap1 (figure 5A-B).

**Figure 5:**
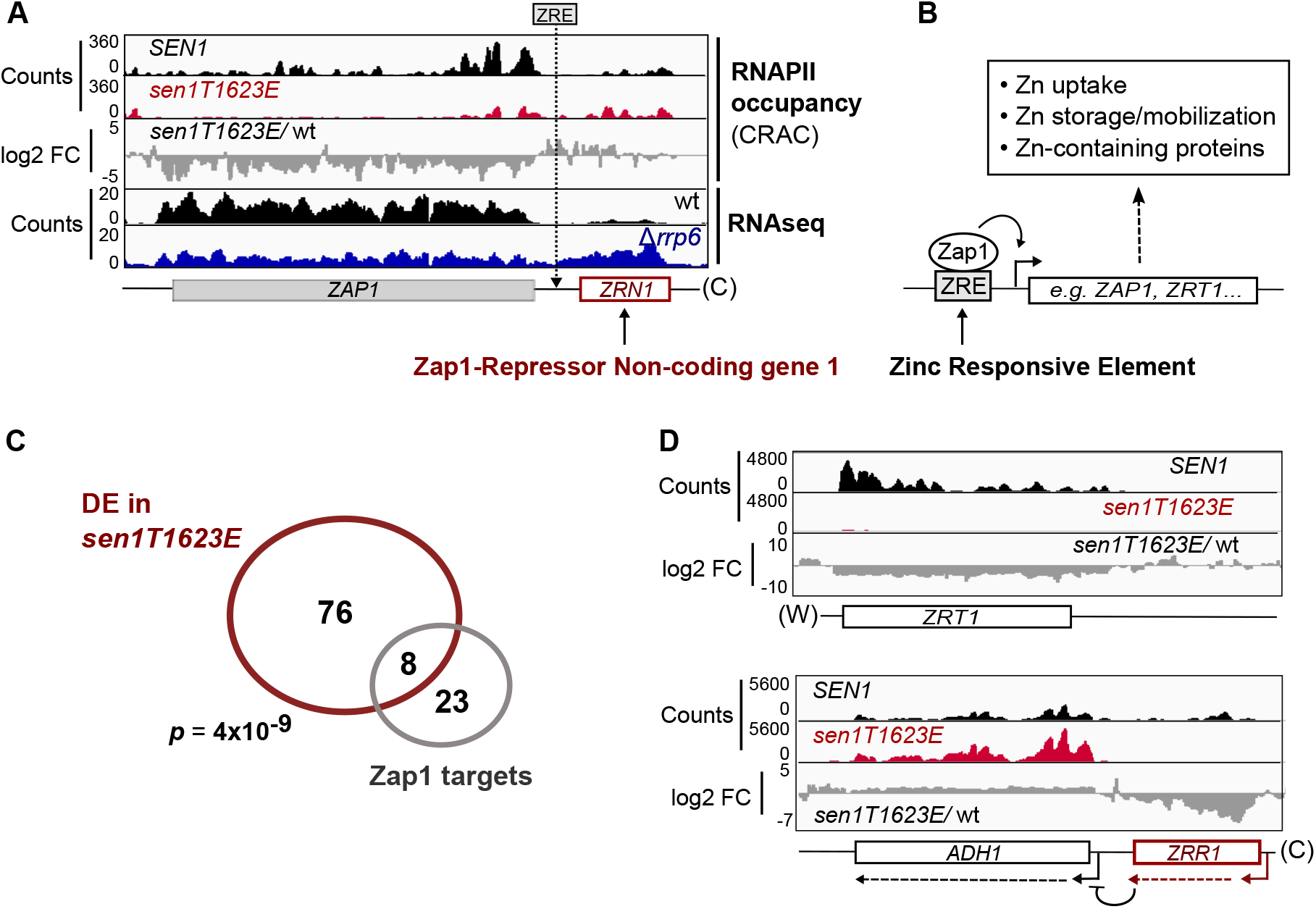
Repression of *ZAP1* underlies a fraction of changes in gene expression provoked by Sen1 T1623E mutation. **A)** IGV screenshot showing the *ZAP1* gene and the upstream non-coding gene *ZRN1*. The location of the previously identified ZRE at the *ZRN1-ZAP1* intergenic region is indicated. The RNAseq datasets will be published elsewhere. **B)** Schematic of the mode of action of Zap1 at regulated promoters and summary of the main processes regulated by Zap1. **C)** Venn diagram showing the overlap between the set of DE genes and Zap1 direct regulatory targets. *p* represents the associated p-value calculated with the Fisher’s exact test. **D)** Examples of well-characterized Zap1-targets that are deregulated in the *sen1T1623E* mutant. Expression of the *ZRR1* induces repression of *ADH1* by transcriptional interference.

### Phosphorylation of Sen1 T1623 partially mimics conditions of Zn excess

Repression of *ZAP1* expression could be responsible for at least part of the changes in gene expression provoked by the Sen1 T1623E mutation. Indeed, we observed a significant overlap between the set of DE genes and previously described Zap1 direct targets (34, 35). Furthermore, the expression pattern of Zap1 target genes in the *sen1T1623E* mutant is consistent with decreased levels of the Zap1 regulator. We observed, for instance, strong repression of *ZRT1*, which encodes a high-affinity Zn permease and is typically activated by Zap1, as well as repression of the non-coding gene *ZRR1* and concomitant upregulation of *ADH1*, which is under the control of *ZRR1* (figure 5D and S5). The fact that we observe a slight induction of some genes typically activated in response to Zn deficiency (e.g. *ZRT1* and *ZRR1*) indicates that in the growth conditions we have employed in our analyses (i.e. growth in minimal medium) cells experience a moderate Zn limitation, consistent with former transcriptome analyses performed in the same conditions (36).

We next aimed at understanding to what extent the deregulation of genes involved in Zn homeostasis could be linked to the observed changes in expression in genes belonging to the other functional categories that we have defined (e.g. mitochondrial function, cell wall and membrane related functions, etc; figure 4D). To address this question, we compared the expression pattern of our DE genes in the *sen1T1623E* mutant relative to the wt, in *Δzap1* relative to wt and in Zn excess, where *ZAP1* is naturally repressed, relative to Zn limitation using available microarray datasets (34). An important limitation of these analyses is the different transcriptional readout provided by the datasets employed. While our CRAC data inform about the levels of nascent transcription, the microarray data provide the total transcript amount, and therefore accumulate both transcriptional and post-transcriptional levels of regulation. Notwithstanding this limitation, all the genes directly involved in Zn homeostasis presented the same trend in the three datasets, as expected (figure 6A). Strikingly, we observed that a large fraction of the remaining genes responded in a very similar way to the Sen1 T1623E mutation and the *ZAP1* deletion. This suggests that the main function of Sen1 phosphorylation at T1623E could be to repress *ZAP1* expression. Notably, this observation concerned the vast majority of genes involved in mitochondrial function, which appeared mainly repressed. Many genes encoding components of the mitochondrial respiratory chain were downregulated in the two mutants as well as in wt cells under conditions of Zn excess, suggesting that under Zn excess repression of *ZAP1* expression could directly or indirectly promote a reduction of the respiratory activity (see discussion for possible explanations). However, we observed that at least one gene encoding a component of the complex V of the electron transport chain, *ATP16*, is located just downstream of an annotated non-coding gene that depends on the NNS-complex for transcription termination (CUT060, see figure S4D). A former study has shown that mutation of the Nrd1 component of the NNS-complex impairs transcription termination at CUT060 and the repression of *ATP16* expression (37). Similarly, in the *sen1T1623E* phospho-mimetic mutant we observed a moderate increase in the RNAPII signal at the CUT060*-ATP16* intergenic region suggesting a termination defect at CUT060 that could underlie the decrease in *ATP16* transcription observed in the mutant (figure S4D).

**Figure 6:**
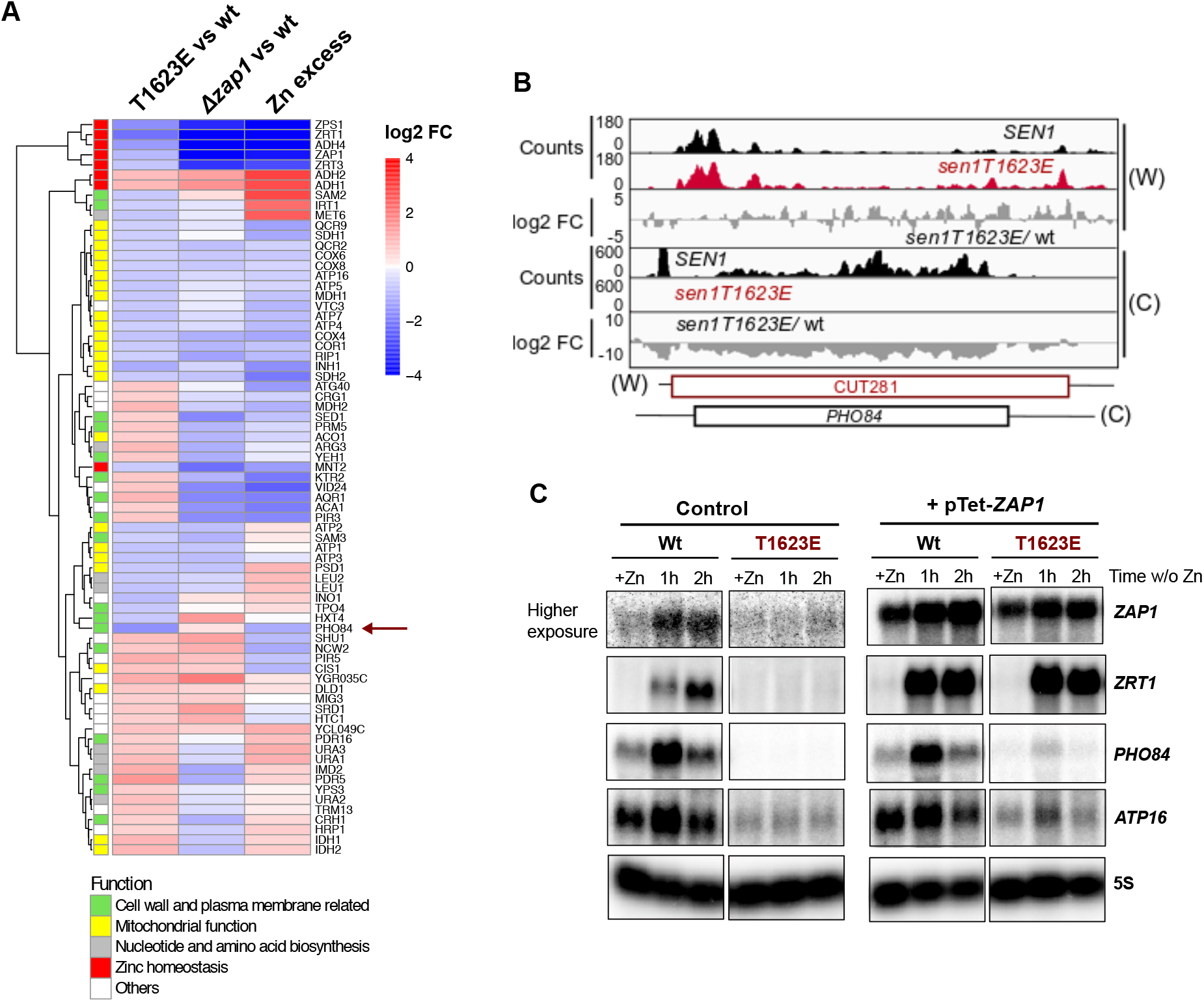
The Sen1 T1623E mutation partially mimics conditions of zinc excess. **A)** Heatmap and hierarchical clustering analyses of the expression profile of DE genes in the indicated datasets. The “*¹¹zap1* vs wt” dataset correspond to conditions of Zn starvation. “Zn excess” is an abbreviation of Zn excess vs limitation. Genes were colored according to the functional category that was assigned to them using the same color code as in figure 4E. **B)** IGV screenshot of CRAC datasets showing the *PHO84* gene and the antisense repressor non-coding gene. **C)** Northern blot analyses of the expression of various genes deregulated in the *sen1T1623E* mutant in conditions of Zn excess (+Zn) or limitation (w/o Zn). Zn excess was reached by growing cells in minimal medium in the presence of 3 mM ZnCl_2_, whereas Zn starvation was induced by shifting exponentially growing cultures to minimal medium with 1 mM EDTA for the indicated times. Representative blot of one out of two independent experiments that yielded similar results. All samples were migrated in the same gel, transferred to the same blot and hybridized in parallel with the same probe except for the probe detecting *ZAP1* mRNA. The image of *ZAP1* RNA detection in control samples (i.e. harbouring an empty vector) correspond to a much higher exposure than the pTet-*ZAP1* samples (i.e. harbouring a centromeric plasmid overexpressing *ZAP1* from a tetracyclin repressible promoter) because endogenous *ZAP1* RNA levels are very low.

Strikingly, we observed a group of genes whose expression pattern was similar in the *sen1T1623E* and Zn-excess datasets but not in the *¹¹zap1* dataset. These include for instance genes involved in detoxification (e.g. *PDR16* and *PDR5*) and, most interestingly, *PHO84*. The latter is probably the best-characterized example of gene repressed by an antisense non-coding gene dependent on the NNS-pathway. Former reports showed that mutations impairing the function of the NNS-complex provoke increased transcription antisense to the *PHO84* promoter and, consequently, repression of *PHO84* expression (16, 38). We observed the same effect in the *sen1T1623E* mutant, strongly suggesting that the phosphorylation at T1623 induces *PHO84* downregulation (figure 6B). Similar repression levels were observed under conditions of Zn excess. *PHO84* encodes a high-affinity phosphate permease, however previous studies showed that this protein can also function as a low-affinity Zn transporter (39) and therefore, its expression might be modulated in response to Zn intracellular concentration. To further test whether repression of *PHO84* and *ATP16* in the *sen1T1623E* mutant is independent on *ZAP1* regulation, we analysed the expression of these genes in a wt and a *sen1T1623E* mutant background in the absence and in the presence of a plasmid overexpressing the *ZAP1* gene from an heterologous promoter. To assess the influence of Zn intracellular concentration on the expression of these genes, we performed these experiments under Zn excess and upon 1 or 2h under Zn starvation (figure 6C). Expression of endogenous *ZAP1* was undetectable under Zn excess and clearly induced after 1h of growth in the absence of Zn in the wt. As expected, in the *sen1T1623E* mutant *ZAP1* mRNA levels remained low in all conditions as if this mutation, and therefore the corresponding mimicked phosphorylation, locked *ZAP1* expression in a Zn-excess repressed state. Consistently, expression of the Zap1-regulated *ZRT1* gene remained repressed in the *sen1T1623E* mutant. However, this effect was completely suppressed by overexpression of *ZAP1*, supporting the idea that deregulation of Zap1-dependent genes in the *sen1T1623E* mutant is mediated by the repression of the *ZAP1* gene. Interestingly, unlike the gene encoding the high-affinity Zn transporter Zrt1, whose expression was more strongly induced as Zn limitation become more severe (i.e. after 2h without Zn in the medium), *PHO84* expression levels were strongly increased early after Zn withdrawal from the medium and decreased subsequently, consistent with its proposed role as a low-affinity Zn transporter. Importantly, *PHO84* expression remained strongly repressed in all conditions in the *sen1T1623E* mutant and this repression was not suppressed by overexpression of *ZAP1*, confirming that the effect of the Sen1 T1623E mutation on *PHO84* expression is independent on *ZAP1* repression. The gene *ATP16* exhibited a similar behavior, namely a clear repression in the *sen1T1623E* mutant regardless the expression levels of *ZAP1*, indicating also a Zap1-independent effect on *ATP16* expression. We observed some fluctuation of *ATP16* expression levels over the different growth conditions, possibly reflecting a response of this gene to stress due to Zn excess or limitation.

Taken together these results suggest that the T1623E mutation partially mimics conditions of Zn excess by inducing directly the repression of *ZAP1* and two other genes, *PHO84* and *ATP16*, and, thus, these conditions could be the origin of the T1623 phosphorylation (figure 7).

**Figure 7:**
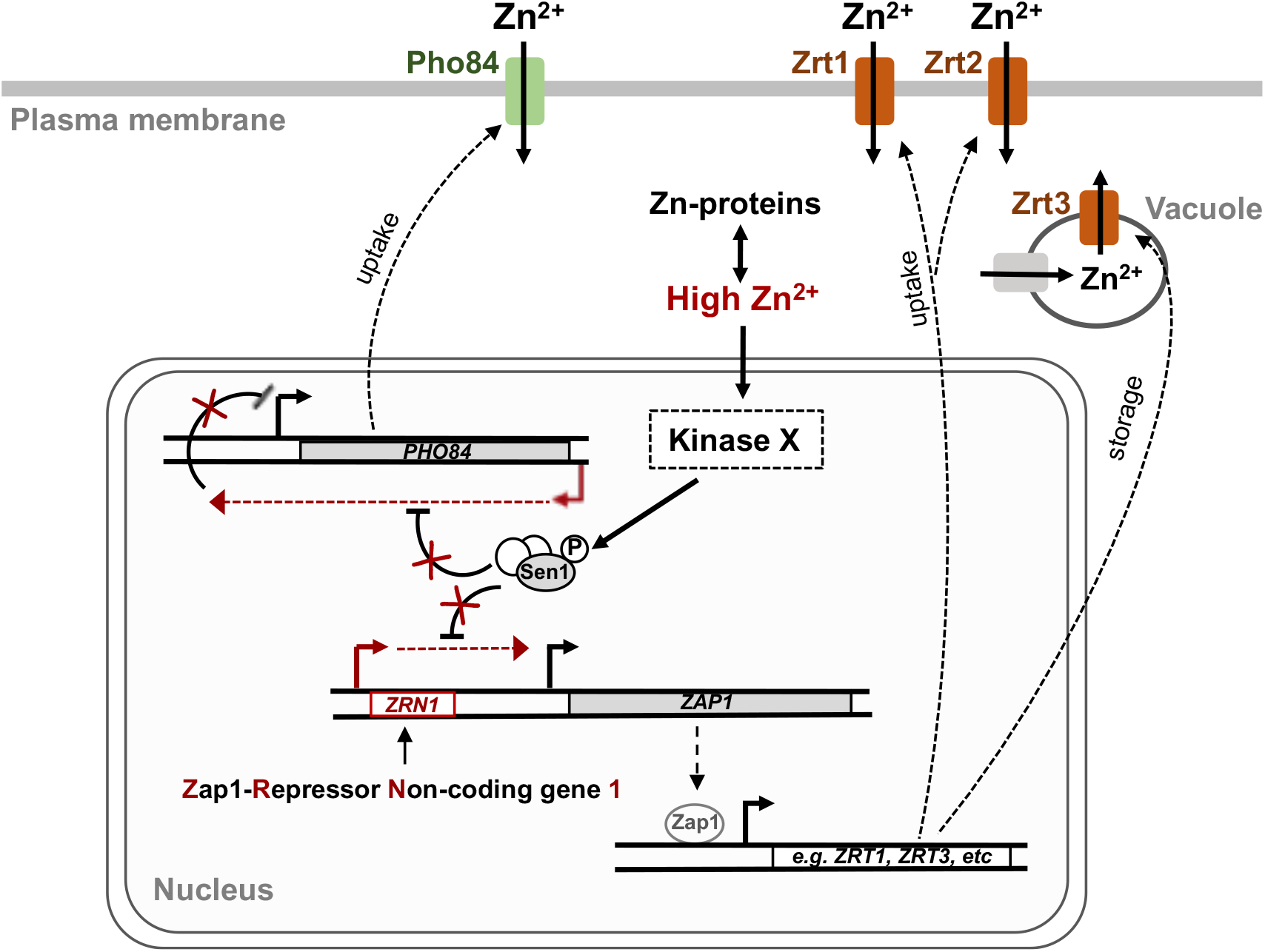
Working model for the role of Sen1 phosphorylation at T1623 in the regulation of gene expression. The NNS-complex normally terminates transcription of *ZRN1* as well as the ncRNA antisense to *PHO84*. Under Zn excess, a kinase yet to be identified would phosphorylate Sen1 to reduce the efficiency of transcription termination and promote repression of *ZAP1* and *PHO84*. Downregulation of *ZAP1* expression would lead to decreased levels of Zap1 protein and, consequently, the regulation of genes involved in Zn homeostasis, as for instance genes responsible for Zn uptake and storage and genes encoding Zn-containing proteins. Repression of *PHO84* expression, encoding a low-affinity Zn transporter, would also contribute to the regulation of Zn homeostasis.

## DISCUSION

Non-coding transcription plays a major role in the regulation of gene expression. In metazoans ncRNAs exist in multiple flavors and can employ a variety of transcriptional and post-transcriptional mechanisms to modulate gene expression (reviewed in 40). However, the transcription of ncRNAs itself can also promote gene regulation, typically by a mechanism of transcriptional interference (reviewed in 41). Budding yeast employs almost exclusively this regulatory strategy. In most cases of repression by transcriptional interference, the non-coding gene is expressed only under the specific condition where repression of a downstream or antisense gene is required (reviewed in 42). However, under all conditions studied, non-coding transcription is widespread and the genome of *S. cerevisiae* very compact, so a large number of genes are transcribed just downstream of or antisense to non-coding genes (::22% of genes, see figure 4) and could, therefore, be potentially subjected to transcriptional interference. Undesired gene repression is prevented by the NNS-complex, which induces early termination at most non-coding transcription units in yeast. Indeed, nuclear depletion of the Nrd1 component of the NNS complex was proposed to cause directly deregulation of about 300 genes by transcriptional interference as well as multiple additional indirect effects in gene expression (4, 17). This highlights the important role of the NNS-complex as a safeguard of genome expression but also raises the question whether the activity of the NNS-complex could be modulated under particular conditions to promote the regulation of specific genes.

Former studies have unveiled mechanisms that could regulate the action of the NNS-complex. For instance, it has been shown that several stress-responsive genes can be repressed by premature termination by the NNS-complex and alternative TSS usage under stress conditions allows excluding Nrd1 and Nab3 recognition sequences from the 5’ UTR and, thus, escaping NNS binding and transcription termination (43). In addition, several phosphorylation sites have been identified in Nrd1 (44) and very recently several methylation sites have been detected in both Nrd1 and Nab3 (45), but a role for these modifications in the regulation of transcription termination has not been demonstrated. The levels of the Sen1 protein have shown to be modulated during the cell cycle (46). Although the function of this modulation is not completely understood, it has been proposed that it could be important to prevent aberrant termination, possibly by adjusting the amount of termination factor to the actual transcriptional input at each phase of the cell cycle. In the present study we describe the identification and characterization of a phosphorylation at the catalytic domain of Sen1 and demonstrate that this modification modulates negatively the capacity of Sen1 to promote transcription termination. The fact that the phosphorylated residue, a threonine, is evolutionary conserved from yeast to humans together with our evidences showing that a similar but non-phosphorylatable amino acid at the same position can support Sen1 function *in vivo* and *in vitro* led us to propose that a threonine has been maintained in Sen1 proteins for regulatory purposes (figures 1-2).

Despite provoking relatively widespread effects on non-coding transcription termination, a phospho-mimetic mutation in the identified phosphosite, T1623, led to changes in the expression of only a small number of genes. Among them, very few were likely to be directly repressed by an inefficiently terminated neighboring non-coding gene (figure 4), suggesting a rather specific role for this phosphorylation in gene regulation. Indeed, our biochemical and genomic data support the idea that phosphorylation of T1623 modulates negatively the association of Sen1 with the nascent RNA, which induces a delay in transcription termination of up to few hundred base pairs. Such delay in termination can result in repression of cognate genes under particular circumstances, for instance, when there is a gene immediately downstream of the inefficiently terminated non-coding transcription unit or when the non-coding gene is terminated already sub-optimally and a mild dysfunction in the NNS-complex can provoke exceptionally strong termination defects. In such scenario, the specificity of the regulation by T1623 phosphorylation would be provided by the particular properties of the target genes. Alternatively, specificity could be achieved locally, in other words by modifying only Sen1 molecules at particular *loci*. In this respect, a growing number of kinases have been found to associate with the chromatin and to regulate transcription-related processes (47–49). Sen1 could therefore be a target of one or several of these kinases. The identity of the kinase/s phosphorylating the Sen1 residue T1623 will shed light onto this question. Unfortunately, our attempt to identify the protein/s responsible for this phosphorylation using an *in vitro* screening with purified kinases was not successful (data not shown). Different approaches will be needed to accomplish this challenging goal in the future.

Among the genes that were differentially expressed in the presence of the Sen1 T1623E phospho-mimetic mutation, a group of genes involved in Zn homeostasis attracted our attention. Zn is an important protein cofactor. It has been estimated that around 10% of yeast proteins require the association with Zn for their enzymatic activity and/or structural stability (50). However, Zn excess can also be toxic for instance because it can compete with other metal ions for the active sites of enzymes. Therefore, Zn intracellular levels need to be tightly controlled, which involves mainly the regulation of Zn uptake and trafficking and the expression of Zn-associated proteins. As mentioned above, a major actor in this process is the transcriptional activator Zap1, which directly controls the expression of genes encoding Zn transporters (e.g. *ZRT1* and *ZRT2*), Zn-containing proteins as well as many genes involved in the cellular adaptation to Zn limitation (34, 35). The activity of Zap1 is highly regulated by Zn at different levels. As outlined above, Zap1 is a DNA binding protein that recognizes ZRE motifs at the promoter region of its target genes and recruits several coactivators through two different transactivation domains (33). At high concentrations, Zn can interact with several regions of Zap1 and modulate negatively the activity of both the transactivation domains and the DNA-binding domain (reviewed in 51). In addition, Zap1 is transcriptionally autoregulated, in other words, it binds to and activates its own promoter (52). In the present study we describe an additional layer of transcriptional regulation that relies on a non-coding gene located just upstream of the *ZAP1* gene and that we have named *ZRN1* (figure 5). Unlike the other formerly reported non-coding genes involved in the regulation of Zn homeostasis, *ZRR1* and *ZRR2*, whose expression is activated in response to Zn limitation, *ZRN1* seems constitutively expressed and acquires the capacity to repress *ZAP1 via* the negative modulation of its transcription termination upon Sen1 T1623 phosphorylation. The expression pattern of Zn homeostasis genes, as well as others (figure 5 and 6) in the *sen1T1623E* phospho-mimetic mutant suggests that Sen1 would be phosphorylated under Zn excess. Both the amount of active Zap1 proteins, determined by the transcriptional output of the *ZAP1* gene and the intracellular concentration of free Zn, and the affinity of Zap1 for the different versions of its recognition motif found in the regulated promoters determines the Zn-responsiveness of the different Zap1 targets. While some genes are activated under mild Zn limitation, others require severe Zn starvation to be activated (35). The combined action of multiple transcriptional and post-transcriptional levels of regulation on Zap1 could be important to better adjust the expression levels of the different Zap1 targets to the actual cellular needs and avoid wasteful production of proteins (e.g. high-affinity Zn transporters) when they are not required (e.g. under Zn-replete/excess conditions). These different mechanisms of regulation of Zap1 activity likely act in a at least partially redundant manner. However it is possible that in a natural environment, where yeast might be subject to multiple simultaneous nutritional and physical stresses, all these mechanisms are necessary for the accurate regulation of Zn homeostasis genes which could be important for cell fitness.

In addition to the repression of *ZAP1*, the Sen1 T1623E phospho-mimetic mutation induces downregulation of the expression of *PHO84*. Repression of this gene is independent on the effect of the Sen1 mutation on the expression of *ZAP1* (figure 6C). However, *PHO84* seems also repressed under Zn excess (figure 6A and C). This gene encodes a high-affinity phosphate permease that can also function as a low-affinity Zn transporter and its overexpression leads to toxicity due to heavy metal accumulation (39, 53), which might explain the necessity to keep *PHO84* expression low under Zn excess. *PHO84* is one of the first and best-characterized examples of gene repressed by antisense transcription in budding yeast (54). Transcription of the antisense non-coding gene (CUT281) is terminated by the NNS-complex and in different mutant contexts where the activity of the NNS-complex is compromised, impaired termination results in increased antisense transcription through the *PHO84* promoter region and, consequently, the repression of *PHO84* by transcriptional interference (16, 38). Similarly, the phospho-mimetic T1623E mutation reduces the efficiency of transcription termination at CUT281 and leads to an accumulation of RNAPIIs antisense to the promoter of *PHO84* as well as a decrease in *PHO84* transcription. Strikingly, the *PHO5* gene, which shares a similar configuration with an antisense NNS-dependent CUT (16) but whose function is exclusively related to the phosphate metabolism, did not exhibit the same behavior as *PHO84* (data not shown). This supports the idea that the Sen1 T1623 phosphorylation could play a specific role in the regulation of zinc homeostasis or related processes. Therefore we propose that under Zn excess phosphorylation of Sen1 at T1623 induces the repression of *PHO84* expression to further contribute to the regulation of Zn intracellular concentration (figure 7).

The most represented category within the genes that respond to the Sen1 T1623E phospho-mimetic mutation, and, therefore, most likely to the phosphorylation of T1623, is the mitochondrial-related function. Indeed 40% of genes encoding components of the mitochondrial respiratory chain were significantly downregulated in the mutant (figure 4). Strikingly, we observed a similar behavior for many of these genes in the *¹¹zap1* mutant as well as in Zn excess relative to limitation (figure 5). Former data indicate that the excess of Zn can interfere with iron metabolism and provoke an increase in the production of reactive oxygen species or ROS (55, 56). Because several components of the mitochondrial respiratory chain use iron as a cofactor (57) and mitochondria are a major source of ROS production, it is possible that an excess of Zn might affect the stability and function of mitochondrial respiratory complexes. Yeast cells might therefore adapt their energetic metabolism to conditions of Zn excess by reducing their respiratory capacity, through the repression of the components of the respiratory chain, and promoting the use of fermentative pathways for energy production. Indeed, budding yeast can use a mixed fermentative-respiratory metabolism and in the presence of high levels of glucose it employs preferentially fermentation (58). The fact that these genes appeared also repressed in the *¹¹zap1* datasets, which were generated under Zn limitation, suggests that the direct and indirect effects of the lack of Zap1 protein on gene expression might be interpreted by the cell as conditions of Zn excess, where *ZAP1* is naturally repressed. Our results show, however, that at least one gene encoding a mitochondrial component, *ATP16*, might be also directly repressed by Sen1 phosphorylation via a termination defect at an upstream non-coding gene (figure S4D and 6C). This regulation could cooperate with other Zap1-dependent mechanisms to modulate mitochondrial functions in response to Zn conditions.

These data support a model according to which under conditions of Zn excess, phosphorylation of Sen1 T1623 would promote the repression of *ZAP1* and *PHO84*, to regulate the intracellular concentration of Zn, as well as the downregulation of components of the mitochondrial respiratory chain to contribute to the appropriate metabolic adaptations.

Overall, our results expand the functional dimension of pervasive transcription by showing that modulation of the efficiency of non-coding transcription termination can promote the regulation of gene expression.

## MATERIAL AND METHODS

### Protein sequence alignments

The sequence of Sen1 protein from *Saccharomyces cerevisiae* (Q00416) was aligned with that of the orthologues from *Candida albicans* (XP_723150), *Schizosaccharomyces pombe* (Q92355), *Mus musculus* (A2AKX3), zebrafish (E7FBJ2), and *Homo sapiens* (Q7Z333) as well as with Upf1 sequence from *S. cerevisiae* (P30771) using clustal omega (https://www.ebi.ac.uk/Tools/msa/clustalo). Sequence alignment was visualized with the jalview software.

### Protein immunoprecipitation experiments and mass spectrometry analyses

Yeast extracts were prepared by standard methods. Briefly, cell pellets were resuspended in lysis buffer (10 mM sodium phosphate pH 7, 200 mM sodium acetate, 0.25% NP-40, 2 mM EDTA, 1 mM EGTA, 5% glycerol) containing protease inhibitors, frozen in liquid nitrogen and lysed using a Retsch MM301 Ball Mill. Protein extracts were clarified by centrifugation at 13000 rpm for 45 min at 4°C and incubated with RNase A (20 μg/ml) for 20 min at RT. Extracts were then incubated with IgG-coupled magnetic beads (Dynabeads, Thermo Fisher, 30 mg/ml) for 2h at 4°C and beads were extensively washed with lysis buffer. Beads were washed with H_2_O and bound proteins were directly subjected to trypsin digestion in 25 mM NH_4_HCO_3_ over-night at 37°C. The resulting peptides were finally analysed by mass spectrometry using an Orbitrap Fusion equipped with an easy spray ion source and coupled to a nano-LC Proxeon 1200 (Thermo Scientific, Waltham, MA, USA).

### Construction of yeast strains and plasmids

Yeast strains used in this paper are listed in table S5. Gene tagging was performed with standard procedures (59, 60) using plasmids described in table S6. Strains DLY3381 and DLY3340 expressing *sen1T1623V* and *sen1T1623E*, respectively were constructed by transforming a *Δsen1* strain harbouring the *URA3*-containing plasmid pFL38-*SEN1* (DLY2767) with the product of cleavage of pFL39-*sen1T1623V* (pDL970) or pFL39-*sen1T1623E* (pDL921) with restriction enzymes MluI, BstZ17I and Bsu36I. Cells capable of growing on 5-FOA were then screened by the absence of pFL38-*SEN1* and pFL39-derived plasmids and the presence of the desired mutation in the chromosome was verified by DNA sequencing.

Plasmids used in this study are listed in table S6. Plasmids expressing different *SEN1* variants were constructed by homologous recombination in yeast. Briefly, a DNA fragment harbouring the desired mutation was obtained by mutagenic overlapping PCR. Then, a wt yeast strain was transformed with a linearized vector and the corresponding PCR fragment, which contains 40–45 bp sequences at both ends allowing recombination with the vector. Clones were screened by PCR, and the positive ones were verified by sequencing. The same procedure was employed for the construction of plasmid pDL1016 overexpressing ZAP1 from the pTet promoter.

Plasmids for overexpression of the T1623V and T1623E variants of Sen1 HD in *E. coli* were generated by introducing the corresponding mutations in plasmid pDL893 using the QuikChange site-directed mutagenesis kit (Stratagene).

### Northern blot assays

Yeast cells used for northern blot assays were typically grown on the appropriate medium, depending on the experiment, to OD_600_ 0.3 to 0.6. For experiments performed under different Zn concentrations cells were grown in DO-TRP supplemented with 3 mM ZnCl_2_ (i.e. Zn excess) to OD_600_ 0.5 and then transferred to DO-TRP containing 1 mM EDTA for 1h or 2h to induce conditions of Zn starvation. Unless otherwise indicated, cells were grown at 30°C. Cells were harvested by centrifugation and RNAs were prepared using standard methods. Samples were separated by electrophoresis on 1.2% agarose gels, and then transferred to nitrocellulose membranes and UV-crosslinked. Radiolabeled probes were typically generated by random priming of PCR products covering the regions of interest with Megaprime kit (GE Healthcare) in the presence of α-^32^P dCTP (3000 Ci/mmol). Oligonucleotides used to generate the PCR probes are indicated in table S7. Hybridizations were performed using a commercial buffer (Ultrahyb, Ambion) at 42°C and after extensive washes, membranes were analysed by phosphorimaging.

### Protein expression and purification

The different Sen1 HD versions were fused to a C-terminal His_8_-tag coupled to *Vibrio cholerae* MARTX toxin cysteine protease domain (61)and expressed from the T7 promoter (see plasmids in table S6) in *Escherichia coli* strain BL21 (DE3) CodonPlus (Stratagene). Overexpression was induced by growth in auto-inducing medium (62) at 20°C overnight. Protein extracts were prepared by sonication of the resuspended cell pellets and subsequent clarification by centrifugation at 20,000 *g* for 30 min at 4°C. Cells were lysed in binding buffer containing 50 mM Tris-HCl pH 8, 500 mM NaCl, 2 mM MgCl_2_, 10 mM imidazole, 10% (v/v) glycerol, 1 mM DTT, benzonase, and protease inhibitors. The proteins were bound to a Ni-NTA sepharose beads (Qiagen), extensively washed with binding buffer and eluted in buffer containing 50 mM Tris-HCl pH 8, 150 mM NaCl, 2 mM MgCl_2_, 5% glycerol and 1 mM DTT. The eluates were then subjected to heparin affinity chromatography on a HiTrap Heparin HP column (GE Healthcare) using buffer A for binding (50 mM Tris–HCl pH 7.5, 200 mM NaCl, 2 mM MgCl_2_, 5% glycerol and 1 mM DTT) and buffer B for elution (50 mM Tris–HCl pH 7.5, 1 M NaCl, 2 mM MgCl_2_, 5% glycerol and 1 mM DTT). Proteins were dialysed against buffer containing 50 mM Tris-HCl, pH 7.5, 350 mM NaCl, 2 mM MgCl_2,_ 50% glycerol and 1 mM DTT and stored at –80°C.

RNAPII core enzyme was purified from *S. cerevisiae* strain BJ5464 (63) by Ni^2+^-affinity chromatography followed by anion exchange on a MonoQ column basically as previously described (28). Recombinant His_6_-tagged Rpb4/7 heterodimer was purified from *E. coli* by Ni^2+^-affinity chromatography followed by gel filtration as previously described (28).

### Electrophoretic mobility shift assays

Experiments were performed in 10 μl-reactions containing 20 mM Tris-HCl pH 7.5, 100 mM NaCl, 2 mM MgCl_2_, 7.5 μM ZnCl_2_,10% glycerol and1 mM DTT. The RNA substrate (4 nM final concentration) was incubated with increasing concentrations of Sen1 HD variants at either 30°C or 37°C for 15 min. Reactions were loaded onto 5% native polyacrylamide gels and the different RNA species were separated by electrophoresis in 0.5xTBE at 100V for 1h. Gels were scanned directly using a Typhoon scanner (GE Healthcare) and analyzed with the ImageQuant TL software. Data were fit using a non-linear regression model and the Kd (dissociation constant) for each protein and condition was calculated using the Prism software (version 9.0.0).

### ATP hydrolysis assays

ATPase assays were performed essentially as previously described (28). Briefly, 10 nM of purified Sen1 proteins was assayed at 30°C or 37°C in 10-μl reactions containing 10 mM Tris– HCl pH 7.5, 75 mM NaCl, 1 mM MgCl_2_, 1 mM DTT, 15% glycerol, and 0.5 ng/μl polyU. The reaction started with the addition of a 250 μM ATP solution containing 0.25 μM of 3000 Ci/mmol α^32^P-ATP as the final concentrations. Aliquots were taken at different time points, mixed with 2.5 volumes of quench buffer containing 10 mM EDTA and 0.5% SDS, and then subjected to thin-layer chromatography on PEI cellulose plates (Merck) in 0.35 M potassium phosphate (pH 7.5). The different radioactive species were analyzed by phosphorimaging.

### *In vitro* transcription termination assays

Termination assays were performed basically as previously described (28). Briefly, ternary ECs were assembled in a promoter-independent manner by first annealing a fluorescently labeled RNA (oligo DL2492, see table S7) with the template DNA (oligo DL3352, see table S7) and subsequently incubating the RNA:DNA hybrid with purified RNAPII. Next, the non-template (NT) strand (oligo DL3353, see table S7) and recombinant Rpb4/7 heterodimer were sequentially added to the mixture. The ternary ECs were then immobilized on streptavidin beads (Dynabeads MyOne Streptavidin T1 from Invitrogen) and washed with transcription buffer (TB) containing 20 mM Tris–HCl pH 7.5, 100 mM NaCl, 8 mM MgCl_2_, 10 μM ZnCl_2_, 10% glycerol, and 2 mM DTT; then with TB containing 0.1% Triton, TB containing 0.5 M NaCl, and finally TB. The termination reactions were performed at 30°C or 37°C in TB in a final volume of 20 μl in the absence or in the presence of 50 nM of different versions of Sen1 HD. Transcription was initiated after addition of a mixture of ATP, UTP, and CTP (1 mM each as the final concentration in the reaction) to allow transcription through the G-less cassette up to the first G of a G-stretch in the NT strand. The reactions were allowed for 15 min and then stopped by the addition of 1 μl of 0.5 M EDTA. After separation of beads and supernatant fractions, beads fractions were resuspended in 8 μl of loading buffer (1× Tris-borate-EDTA, 8 M urea) and boiled for 5 min at 95°C. Transcripts in the supernatant fractions were ethanol precipitated and resuspended in 8 μl of loading buffer. Transcripts were separated by 10% (w/v) denaturing PAGE (8 M urea), and gels were scanned with a Typhoon scanner (GE Healthcare) and analyzed with the ImageQuant TL software (GE Healthcare).

### UV crosslinking and analysis of cDNA (CRAC)

For RNAPII CRAC experiments we employed the procedure reported in Granneman *et al* (29) with several modifications described in Candelli *et al* (64). We used strains expressing an His6-TEV-Protein A-tagged version of the Rpb1 subunit of RNAPII at the endogenous locus and either the wt or the T1623E version of Sen1 (see table S5). Briefly, 2 l of yeast cells were grown at 30°C to OD_600_ ∼ 0.6 in CSM-TRP medium and crosslinked for 50 s using a W5 UV crosslinking unit (UVO3 Ltd). Cells were then harvested by centrifugation at 12,000 *g* for 10 min and the pellets were washed with cold 1× PBS and resuspended in 2.4 ml of TN150 buffer (50 mM Tris pH 7.8, 150 mM NaCl, 0.1% NP-40 and 5 mM β-mercaptoethanol) containing protease inhibitors (Complete EDTA-free Protease Inhibitor Cocktail) per gram of cells. Suspensions were flash-frozen in droplets and cells were lysed using a Ball Mill MM 400 (five cycles of 3 min at 20 Hz). The mixtures were incubated with 165 units of DNase I (NEB) for 1 h at 25°C and then clarified by centrifugation at 20,000 *g* for 20 min at 4°C.

Protein extracts were subjected to IgG affinity purification on M-280 tosylactivated dynabeads coupled with rabbit IgGs (15 μg of beads per sample). After extensive washes with TN1000 buffer (50 mM Tris pH 7.8, 1 M NaCl, 0.1% NP-40 and 5 mM β-mercaptoethanol), the protein– RNA complexes were eluted by digestion with the TEV protease and treated with 0.2 U of RNase cocktail (RNace-IT, Agilent) to reduce the size of the nascent RNA (note that the 3′ end of nascent transcripts is protected from degradation). The eluates were mixed with guanidine– HCl to a final concentration of 6 M and incubated with Ni-NTA sepharose (Qiagen, 100 μl of slurry per sample) o/n at 4°C. After washing beads, sequencing adaptors were ligated to the RNA molecules as described in the original procedure. Protein–RNA complexes were eluted with 400 μl of elution buffer (50 mM Tris pH 7.8, 50 mM NaCl, 150 mM imidazole, 0.1% NP-40, 5 mM β-mercaptoethanol) and concentrated using Vivacon ultrafiltration spin columns. Then, proteins were fractionated using a Gel Elution Liquid Fraction Entrapment Electrophoresis (GelFree) system (Expedeon) following manufacturer’s specifications and the different fractions were monitored for the presence of Rpb1 by SDS–PAGE. The fractions of interest were treated with 100 μg of proteinase K, and RNAs were purified and reverse-transcribed using reverse transcriptase Superscript IV (Invitrogen).

The cDNAs were amplified by PCR using LA Taq polymerase (Takara), and then, the PCR reactions were treated with 200 U/ml of Exonuclease I (NEB) for 1 h at 37°C. Finally, the DNA was purified using NucleoSpin columns (Macherey-Nagel) and sequenced on a NextSeq 500 Illumina sequencer.

### Data processing

CRAC reads were demultiplexed using the pyBarcodeFilter script from the pyCRACutility suite (65). The 5′ adaptor was clipped with Cutadapt and the resulting insert was quality-trimmed from the 3′ end using Trimmomatic rolling mean clipping (66). We used the pyCRAC script pyFastqDuplicateRemover to collapse PCR duplicates using a six-nucleotide random tag included in the 3′ adaptor. The resulting sequences were reverse complemented with the Fastx reverse complement that is part of the fastx toolkit (http://hannonlab.cshl.edu/fastx_toolkit/) and mapped to the R64 genome with bowtie2 using “-N 1”option. Coverage files were normalized to 10^7^ counts.

### Bioinformatic and statistical analyses

Bioinformatic analyses were mainly performed using the Galaxy (http://galaxy.sb-roscoff.fr) and the R frameworks. For metagene analyses, we used the deepTools2 package (67). Strand-specific coverage bigwig files were used as inputs for the computeMatrix tool together with separate annotations for each strand and for each feature (CUTs or snoRNAs), using a bin size of 10 and the TTS as the reference point. Matrices generated for each strand were subsequently combined using the rbind option of the computeMatrixOperations tool and used as the input for the plotProfile tool. We typically represented the median instead of the mean values to minimize the bias towards the very highly expressed features. The log2 FC of the RNAPII signal in the *sen1T1623E* mutant relative to the wt was calculated using the tool bigwigCompare. As for metagene analyses, we generated separate matrices for each strand with the log2 FC bigwig files and the computeMatrix tool, and we subsequently combined them using the computeMatrixOperations tool. Heatmaps were obtained using these matrices as the input for the plotHeatmap tool. For metagene and heatmap analyses we employed the refined CUTs and snoRNA annotations reported in Han et al (14).

Differential expression analyses were performed by counting the reads from two independent biological replicates that mapped at protein-coding genes using the htseq-count tool (68) with the union mode. The count files thus generated were used as input files for the SARTools DESeq2 package (69) with default parameters.

For the analysis of non-coding transcription relative to protein-coding genes we used the CUTs and SUTs annotations reported in Xu et al, (5) and the mRNAs annotations in Challal et al (32). We employed the window bed tool of the bedtools package (70) to detect the genes that overlap with an antisense CUT or SUT or that initiate within a 500-bp window downstream of one of these non-coding genes. When a gene was located both antisense to and downstream of a CUT or SUT we assigned a value of 0.5 gene to each category. To calculate the statistical significance of the enrichment of one particular category of genes (e.g. genes located just downstream of annotated non-coding genes) we performed a Fisher’s exact test with R.

Gene ontology analyses were performed with the Genome Ontology Term Finder tool of the Saccharomyces Genome Database (https://www.yeastgenome.org/goTermFinder).

Heatmap analyses of the expression pattern of DE genes in different backgrounds and conditions were performed with the pheatmap function of R. As input files we used the results of our differential expression analyses together with microarray datasets reported in (34).

## DATA AVAILABILITY

The RNAPII CRAC data have been deposited in NCBI’s Gene Expression Omnibus (GEO) and are accessible through GEO Series accession number GSE174623.

## ACKNOWLEDGEMENT

We thank G. Wentzinger for technical assistance and B. Palancade, D. Libri and the other members of the Libri lab for fruitful discussions. We thank the Roscoff Bioinformatics platform ABiMS (http://abims.sb-roscoff.fr) for providing computational resources and support. This work has benefited from the facilities and expertise of the high throughput sequencing core facility of I2BC (http://www.i2bc.paris-saclay.fr/). We thank the proteomics facility of the Institut Jacques Monod, supported by the Region Ile-de-France (SESAME), Université de Paris and the CNRS, for their technical assistance. We thank C. Sole and F. Posas for their attempt to identify the kinases phosphorylating Sen1 helicase domain.

