## Supplementary material for "Modulated termination of non-coding transcription partakes in the regulation of gene expression"

Haidara, N and O. Porrua

**List of supplementary material:**

-Supplementary figures S1-5.

-Tables S5, S6 and S7.

**Material provided as separate files:**

**-Table S1:** mass spectrometry analyses of Sen1-TAP.

**-Table S2:** results of differential expression analyses in the *sen1T1623E* relative to the wt strain.

**-Table S3:** List of GO terms overrepresented in the set of genes upregulated in the *sen1T1623E* mutant compared to the wt.

**-Table S4:** List of GO terms overrepresented in the set of genes downregulated in the *sen1T1623E* mutant compared to the wt.

### SUPPLEMENTARY FIGURES

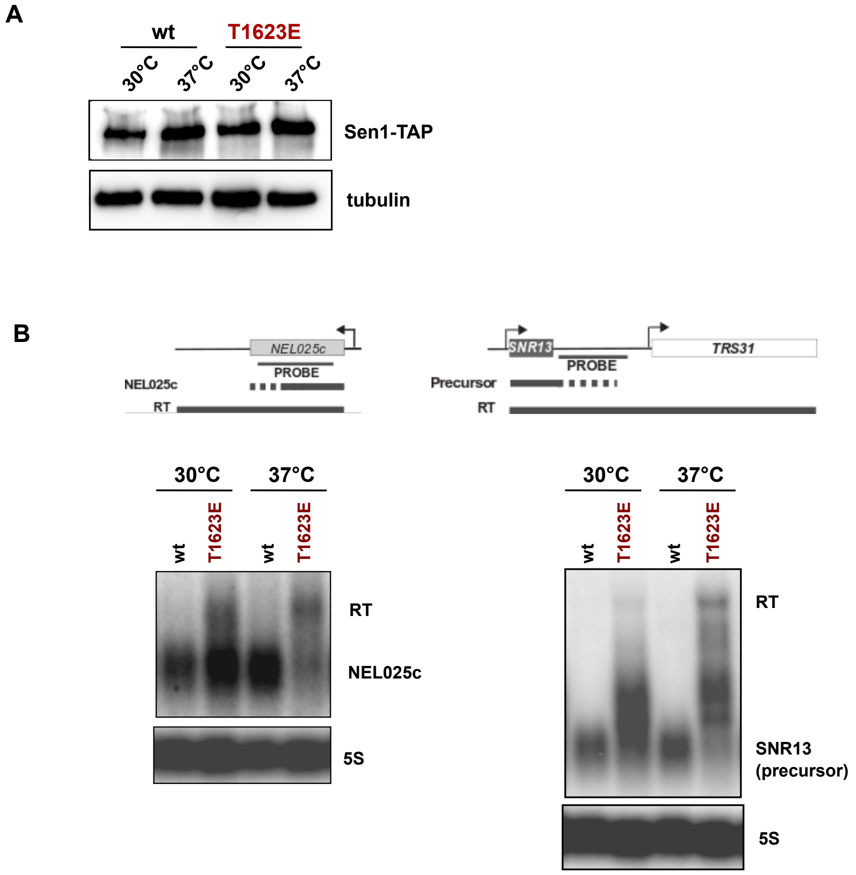


**Figure S1: Phenotypic characterization of the *sen1T1623E* mutant at 37°C.**
**A)** SDS-PAGE analysis of C-terminally TAP-tagged Sen1 wt or T1623E expressed in yeast. Cells were grown at permissive temperature (30°C) and kept at 30°C or at high (37°C) temperature for 2h before being harvested for protein extract preparation. Proteins were detected by western blot using an antibody against the protein A moiety of the TAP-tag. Tubulin is detected as a loading control. Representative gel of one out of three independent experiments with similar results. **B)** Northern blot analysis of ncRNAs typically targeted by Sen1 for transcription termination. Cells were grown at permissive temperature (30°C) or incubated at high (37°C) temperature for 2h. Experiments were performed in a *Δrrp6* background. The 5S RNA is used as a loading control. Probes used for RNA detection are described in table S7.

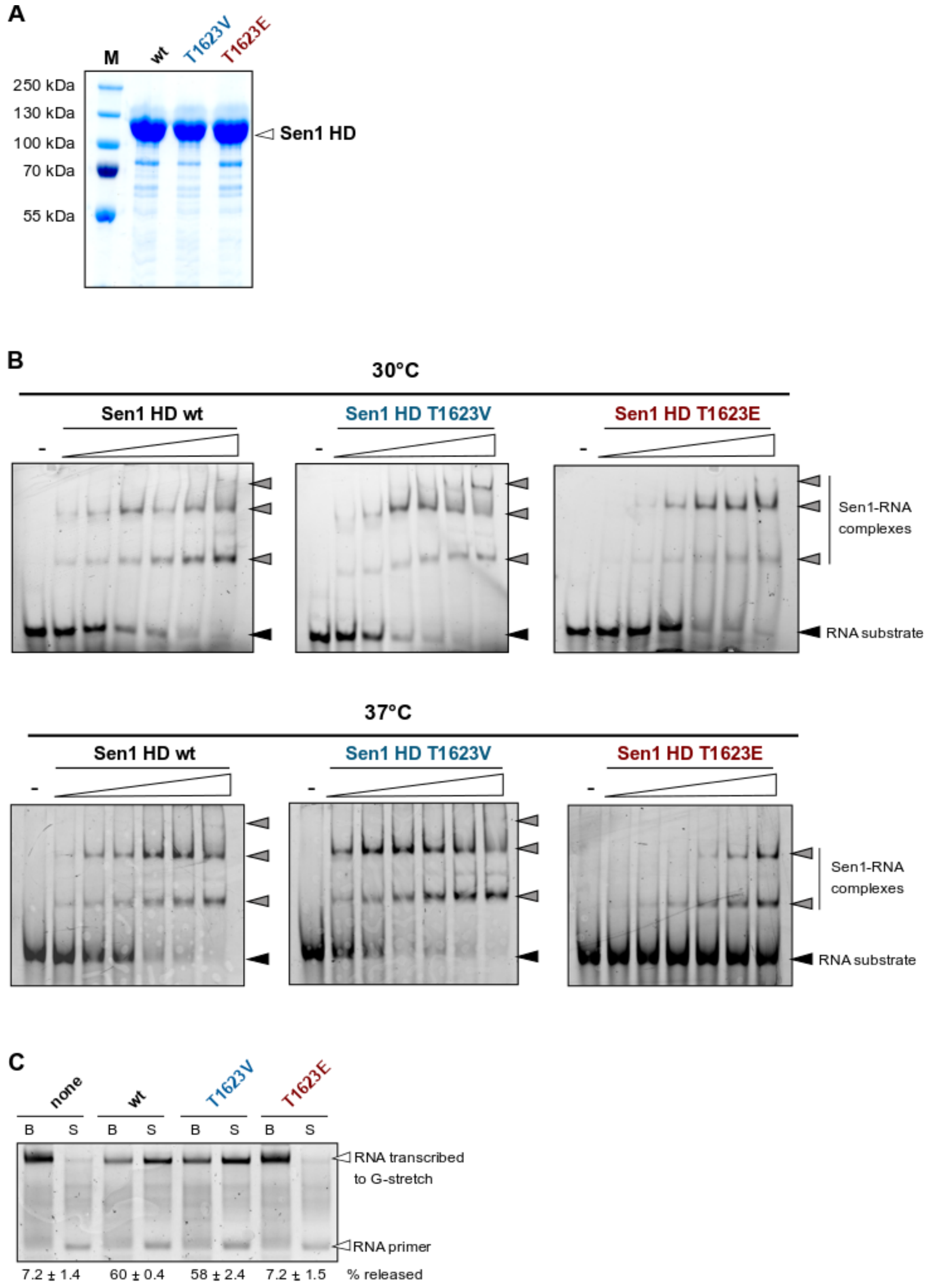


**Figure S2: Related to figure 2.**
**A)** SDS-PAGE analysis of purified Sen1 HD variants used in the experiments in figure 2.
**B)** Representative gels of EMSA summarized in figure 2C. For each experiment, the gels used for the wt, the T1623V and the T1623E mutant version of Sen1 HD were migrated and processed in parallel. A black arrowhead denotes the RNA substrate whereas the Sen1-RNA complexes are indicated by a grey arrowhead. Bands corresponding to several Sen1 HD molecules per RNA can be observed as the protein concentration increases. **C)** Representative gel of IVTT assays performed at 37°C. The values on the bottom of the gel correspond to the average and SD of three independent experiments.

A

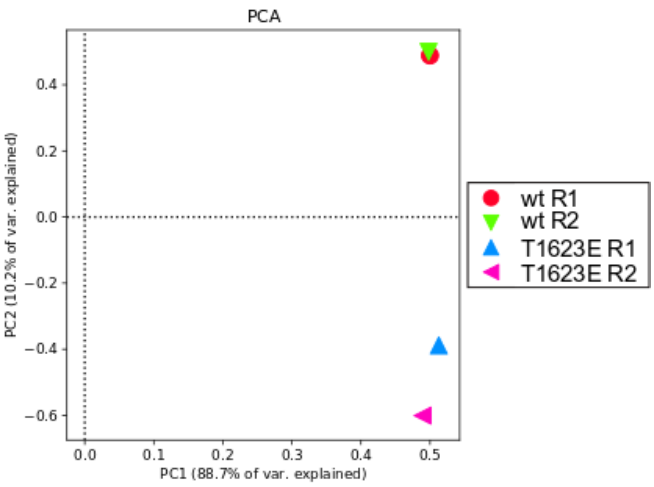


B

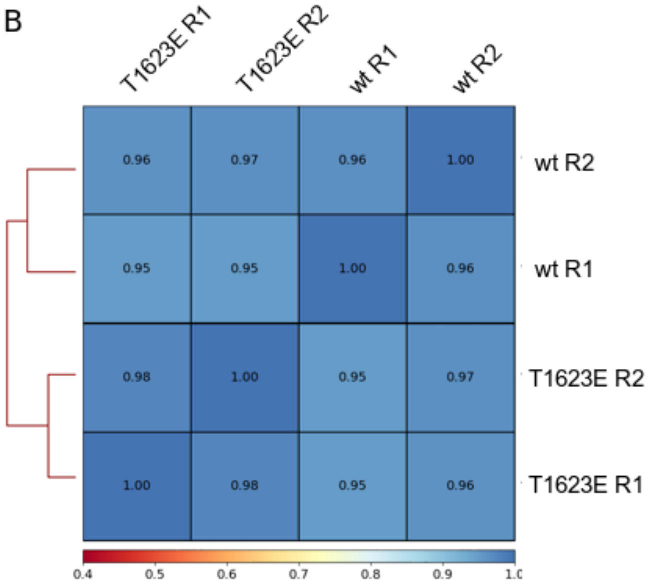


**Figure S3: Principal component analysis and correlation plots illustrating the high reproducibility of the two biological replicates of CRAC experiments in figure 3.**
**A)** Principal component analysis (PCA) performed using the normalized reads coverage on the Watson strand (only the top 1000 bins of 500 bp). R1 and R2 denote the biological replicate 1 and 2, respectively. **B)** Heatmap representation of the correlation between the different datasets calculated as for the PCA with the coverage values of 500 bp bins in the Watson strand. The heatmap values correspond to the spearman coefficient.

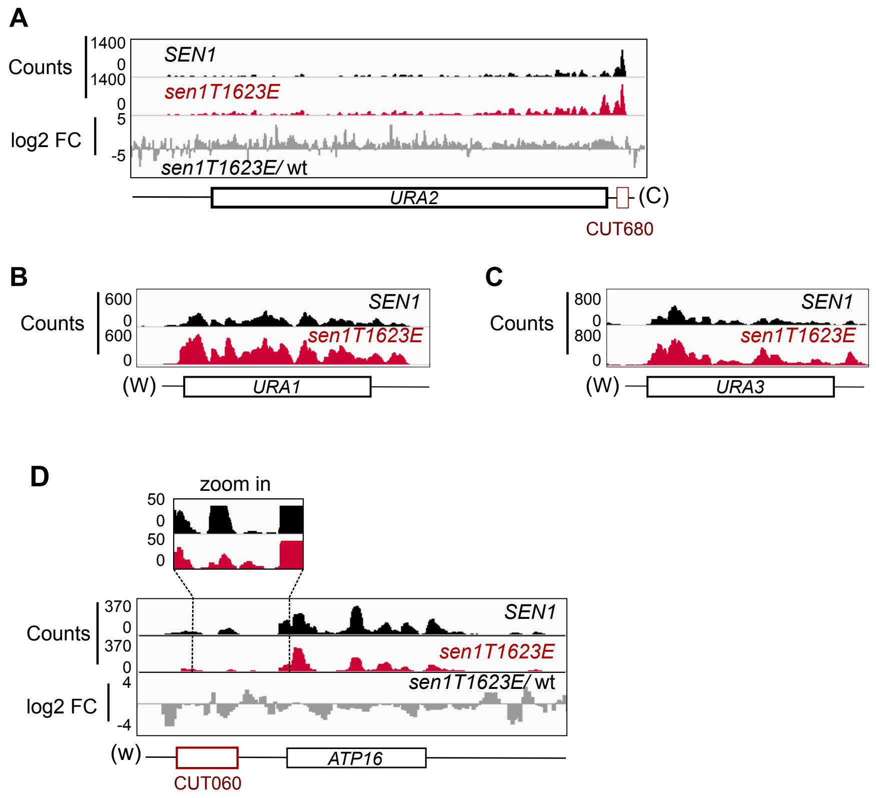


**Figure S4: Deregulation of genes involved in UMP biosynthesis and mitochondrial function by the Sen1 T1623E mutation.**

**A)** IGV screenshot of the *URA2* gene, likely deregulated by inefficient transcription termination at an upstream CUT. **B**) and **C**) IGV screenshots of additional genes of the UMP biosynthetic pathway that are most likely indirectly deregulated by the Sen1 T1623E mutation.

**D**) IGV screenshot of the *ATP16* gene encoding a component of the mitochondrial respiratory chain. Top: Zoom in view of the CUT060-*ATP16* intergenic region to better visualize a moderate increase in the RNAPII signal at this region in the *sen1T1623E* mutant.

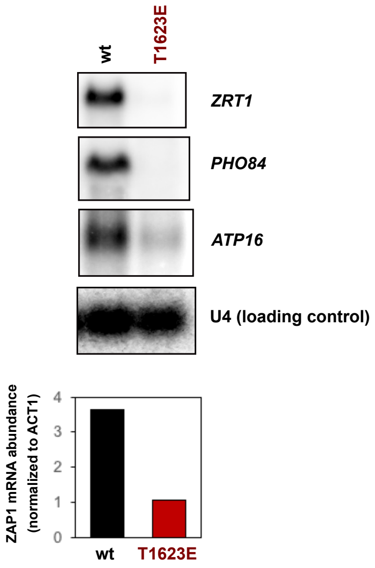


**Figure S5: Northern blot analysis of mRNAs deregulated by the Sen1 T1623E mutation.** RNA samples were prepared from aliquots of the same cultures used for CRAC experiments. Blot corresponding to one of the two biological replicates. The U4 RNA is used as a loading control. Probes used for RNA detection are described in table S7. Because of problems with the detection of *ZAP1* mRNA by northern blot in this experiment, we show the quantification of this RNA by RT-qPCR. Values correspond to the RNA abundance relative to the *ACT1* mRNA and multiplied by 1000.

**SUPPLEMENTARY TABLES**

**Table S5:** List of yeast strains used in this study.

| **Number** | **Name** | **Genotype** | **Source** |
|---|---|---|---|
| DLY671 | BMA | *as W303, Δtrp1* | F. Lacroute |
| DLY814 | *Δrrp6* | *as W303, rrp6::KAN* | (Porrua et al., 2012) |
| DLY2571 | *rpb1-HTP* | *as BMA, rpb1::HTP::TRP1kl* | (Candelli et al, 2018) |
| DLY2767 | *Δsen1/*pFL38-*SEN1* | *as BMA, sen1::KAN, harbouring plasmid pFL38-SEN1* | (Han et al, 2020) |
| DLY3411 | *sen1T1623E, rpb1-HTP* | *as BMA, sen1T1623E, rpb1-HTP:: TRP* | This work |
| DLY3340 | *sen1T1623E* | *as BMA, sen1T1623E* | This work |
| DLY3345 | *sen1T1623E, Δrrp6* | *sen1T1623E, rrp6::KAN* | This work |
| DLY3381 | *sen1T1623V* | *as BMA, sen1T1623V* | This work |
| DLY3389 | *sen1T1623V, Δrrp6* | *as BMA, SEN1::sen1T1623V, rrp6::KAN* | This work |
| DLY2131 | *sen1-TAP* | *as BMA,SEN1::TAP::HIS* | This work |
| DLY3284 | *sen1T1623E -TAP* | *as BMA, sen1T1623E::TAP::URA* | This work |

**Table S6:** List of plasmids used in this study.

| **Name** | **Description** | Source |
|---|---|---|
| pDL693 | $Ap^r$; *ori*ColE1; derivative of pFL39 (TRP) bearing yeast *SEN1* | F. Lacroute |
| pDL772 | $Ap^r$; *ori*ColE1; derivative of pFL38 (URA) bearing yeast *SEN1* | F. Lacroute |
| pDL893 | $Ap^r$; *ori*ColE1, vector for overexpression of proteins from the T7 bearing sen1-HD(1095-1904)CPD-$His_8$ | (Leonaite et al, 2017) |
| pDL921 | $Ap^r$; *ori*ColE1; derivative of pFL39 bearing *sen1-T1623E* | This work |
| pDL926 | $Ap^r$; *ori*ColE1, , vector for overexpression of proteins from the T7 bearing *sen1-HD-T1623E-CPD-* $His_8$ | This work |
| pDL940 | $Ap^r$; *ori*ColE1,, vector for overexpression of proteins from the T7 bearing *sen1-HD-T1623V-CPD-* $His_8$ | This work |
| pDL970 | $Ap^r$; *ori*ColE1; derivative of pFL39 bearing *sen1-T1623V* | This work |
| pDL1016 | $Ap^r$; *ori*ColE1; derivative of pCM185 (TRP) expressing *ZAP1* from a pTet (Tet-Off) promoter. | This work |

**Table S7:** List of oligonucleotides used in this study.

| Name | Sequence (5'-3') | Information/use |
|---|---|---|
| DL1452 | CTTCCCCGTAGAAAATCTTA | Fwd primer to generate by PCR a probe to detect SNR13 |
| DL1120 | TCCGTGTCTCTTGTCCTGCA | Rev primer to generate PCR a probe to detect SNR13 |
| DL474 | GCAAAGATCTGTATGAAAGG | Forward oligo to generate by PCR a probe to detect NEL025C. |
| DL480 | ATCTGACCAGGTCAAGCTAC | Reverse oligo to generate by PCR a probe to detect NEL025C. |
| DL4651 | TGAACCTGCCTGCTAAGTCAGG | Forward oligo to generate by PCR a probe to detect *ATP16* mRNA |
| DL4652 | ATTGCAGCTTCTGCGGCTTC | Reverse oligo to generate by PCR a probe to detect *ATP16* mRNA |
| DL4688 | CGGTGGTGACTACCCACTAT | Forward oligo to generate by PCR a probe to detect *PHO84* mRNA |
| DL4689 | TGGCACCGACCTTACCAGAT | Reverse oligo to generate by PCR a probe to detect *PHO84* mRNA |
| DL2627 | ATTCAAAAGCGAACACCGAATTGACCAT GAGGAGACGGTCTGGTTTAT | Reverse oligo used as a probe to detect U4 snRNA |
| DL377 | ATGTTCCCAGGTATTGCCGA | Forward oligo to generate by PCR a probe to detect *ACT1* mRNA. |
| DL378 | ACACTTGTGGTGAACGATAG | Reverse oligo to generate by PCR a probe to detect *ACT1* mRNA. |
| DL743 | GGTTGCGGCCATATCTACCA | Reverse oligo used as a probe to detect the 5S rRNA |
| DL4659 | GCCATGGGCCCTATGTGTTG | Forward oligo to generate by PCR a probe to detect *ZRT1* mRNA |
| DL4660 | GCGCAGTGTAAGAACCGCTG | Reverse oligo to generate by PCR a probe to detect *ZRT1* mRNA |
| DL2492 | UGCAUUUCGACCAGGC | 5' FAM labeled RNA oligo for performing IVTT assays on immobilized templates |
| DL3352 | CTAGAGGAAACAAACTATAGGAAACGA CCAGGCCCTCAACATCTCTCACCCATC TCCACACGGGGGTTACCCGGCCTGCA | Non-template strand for IVTT assays |
| DL3353 | GGCCGGGTAACCCCCGTGTGGAGATG GGTGAGAGATGTTGAGGGCCTGGTCGT TTCCTATAGTTTGTTTCCT | Template strand for IVTT assays |
| DL3508 | UUCAUUUCAGACCAGCACCCACUCACU ACAACUCACGACCAGGC | 5' FAM labeled RNA to be used as substrate for EMSA experiments |
